# A model screening pipeline for bile acid converting anti-*Clostridioides difficile* bacteria reveals unique biotherapeutic potential of *Peptacetobacter hiranonis*

**DOI:** 10.1101/2021.09.29.462466

**Authors:** Akhil A. Vinithakumari, Belen G. Hernandez, Sudeep Ghimire, Seidu Adams, Caroline Stokes, Isaac Jepsen, Caleb Brezina, Orhan Sahin, Ganwu Li, Chandra Tangudu, Claire Andreasen, Gregory J. Philips, Michael Wannemuehler, Albert E. Jergens, Joy Scaria, Brett Sponseller, Shankumar Mooyottu

## Abstract

*Clostridioides difficile* is an antibiotic-resistant bacterium that causes serious, toxin-mediated enteric disease in humans and animals. Gut dysbiosis and resultant alterations in the intestinal bile acid profile play an important role in the pathogenesis of *C. difficile* infection (CDI). Restoration of the gut microbiota and re-establishment of bacterial bile acid metabolism using fecal microbiota transplantation (FMT) has been established as a promising strategy against this disease, although this method has several limitations. Thus, a more defined and precise microbiota-based approach using bacteria that biotransform primary bile acids into secondary bile acids could effectively overcome these limitations and control CDI. Therefore, a screening pipeline was developed to isolate bile acid converting bacteria from fecal samples. Dogs were selected as a model CDI-resistant microbiota donor for this pipeline, which yielded a novel *Peptacetobacter hiranonis* strain that possesses unique anti-*C. difficile* properties, and both bile acid deconjugation and 7-α dehydroxylating activities to perform bile acid conversion. The screening pipeline included a set of *in vitro* tests along with a precision *in vivo* gut colonization and bile acid conversion test using altered Schadler flora (ASF) colonized mice. In addition, this pipeline also provided essential information on the growth requirements for screening and cultivating the candidate bacterium, its survival in a CDI predisposing environment, and potential pathogenicity. The model pipeline documented here yielded multiple bile acid converting bacteria, including a *P. hiranonis* isolate with unique anti-*C. difficile* biotherapeutic potential, which can be further tested in subsequent preclinical and human clinical trials.

## Introduction

*Clostridioides difficile* infection (CDI) is a serious toxin-mediated enteric disease affecting humans and animals.^1–3^ Antibiotic therapy, gut-dysbiosis, and resultant alterations in the intestinal bile acid profile play important roles in CDI disease pathogenesis.^1, 4^ Annually, approximately half a million people in the United States suffer from CDI, which incurs approximately $5 billion in treatment and prevention costs.^5^ Recent studies indicate that 1 out of every 11 patients aged 65 or older dies within 30 days of CDI diagnosis.^6^ *C difficile* is also a significant animal pathogen that causes serious enteric disease, mortality, and economic losses in a variety of species, especially in pigs with the potential for zoonotic transmission.^7, 8^ Currently, antibiotics are the primary treatment for CDI despite the risk of further aggravation of dysiosis, relapse of infection, and development of antibiotic resistance.^9^ Recently, *C. difficile* has been listed as an urgent threat by the Center for Disease Control and Prevention (CDC) concerning the emergence of antibiotic resistance.^10^ Therefore, the development of an effective, alternative approach for controlling CDI is critical. Restoration of the gut microbiome and re-establishment of normal bile acid metabolism using fecal microbiota transplantation (FMT) has been recently established as a promising and effective strategy against this disease.^11–13^ Unfortunately, the concerns of transmission of bacterial and viral pathogens and reports of deaths associated with FMT highlight the need for more defined and precise microbiome-based approaches for controlling CDI in patients.^14, 15^

The liver primarily synthesizes cholic acid (CA) and chenodeoxycholic acid (CDCA) from cholesterol in hepatocytes. Therefore, CA and CDCA are known as primary bile acids that are conjugated with glycine or taurine to form conjugated primary bile acids (e.g., taurocholic acid (TCA)), which are secreted into the small intestine.^16, 17^ In mice, muricholic acid (MCA) is also synthesized in the liver along with CA and CDCA as a primary bile acid.^18^ Several bile salt hydrolyzing gut bacteria that contain the *bsh* (bile salt hydrolase) gene deconjugate secreted primary bile acids.^17^ At the same time, a few species of 7-α dehydroxylating bacteria, such as *Clostridium scindens* and *Peptacetobacter hiranonis* (previously *Clostridium hiranonis*), that contain the crucial *bai* operon (bile acid-inducible operon), convert these deconjugated primary bile acids into secondary bile acids including deoxycholic acid (DCA) and lithocholic acid (LCA).^16, 17^ Primary bile acid CA facilitates the germination of *C. difficile* spores, whereas secondary bile acids, formed by bacterial conversion of the primary bile acids in the colon strongly inhibit the growth of *C. difficile* in the host.^19–21^ In addition, bile acids directly bind and inhibit the activity of *C. difficile* toxin in the gut and modulate host immune and inflammatory responses.^13, 22, 23^ Long-term antibiotic therapy with resultant dysbiosis in humans precipitate CDI by eliminating bile acid converting bacteria (both *bsh* and *bai* containing species) in the intestine, increasing the available primary bile acids, and reducing the pool of secondary bile acids.^4^ Therefore, bile acid metabolizing bacteria, especially rare *bai* containing species have immense therapeutic value for the treatment of CDI.^19^ As opposed to the current strategy of providing a whole fecal microbiota mixture (FMT) to the patient, a targeted and’ precision microbiome’-based strategy for normalizing the gut bile acid profile has been proposed as a promising alternative for controlling CDI.^19, 24^ Therefore, there is a critical need to identify and isolate *bai* coding bacterial strains and to demonstrate the bile-acid converting ability to exploit their prophylactic and therapeutic potential. These bacteria have immense potential to be used as live biotherapeutic agents to prevent and treat CDI as a functional alternative to antibiotics and conventional FMT or to formulate an enriched or defined FMT preparation.

The presence of bile acid metabolizing bacteria such as *C. scindens* and *P. hiranonis* in the resident gut microbiota is associated with notable resistance to *C. difficile* colonization in different mammalian hosts, especially in dogs.^20, 25^ However, identifying, isolating, and cultivating these unique, invaluable, but fastidious anaerobes is challenging. Therefore, only few isolates of *C. scindens* and virtually none of *P. hiranonis* have been fully characterized in the laboratory for growth conditions, *in vitro* and *in vivo* bile acid conversion ability, gut colonization, and anti-*C. difficile* therapeutic properties. In this study, a model pipeline is described for isolating bile acid converting bacteria, such as *P. hiranonis,* from naturally *C. difficile* resistant hosts. The model takes advantage of dogs as a model host naturally resistant to CDI. This approach involves a series of *in vitro* tests and a novel *in vivo* gut colonization and bile acid conversion model using Altered Schadler Flora (ASF) colonized mice. In addition, this pipeline also provides essential information on the growth requirements of the candidate bacterium (e.g., *P. hiranonis*), its survival in a CDI predisposing gut environment, enterotoxicity, and *in vivo* and *in vitro* bile acid converting abilities. These data are critical for further preclinical infection studies and clinical trials. Thus, this manuscript documents the steps involved in this pipeline that resulted in the identification and characterization of a unique *P. hiranonis* isolate (*P. hiranonis BAI-17*) with anti-*C. difficile* therapeutic value. Additionally, this study establishes *P. hiranonis* as the only bacterium currently known to have both bile acid hydrolyzing and 7-α hydroxylation capabilities, which are demonstrated *in vitro* and *in vivo*. The unique combination of functional *bai* and *bsh* genes within a single bacterial genome makes *P. hiranonis* a promising biotherapeutic candidate for targeted microbiome-based therapy against CDI in humans and animals.

## Results

### Specific *P. hiranonis* strains and other anaerobes from canine feces are selectively isolated in a beta-lactam -fluoroquinolone-bile acid environment

For this study, dogs were selected as the model *C. difficile* resistant donors due to their natural resistance to CDI, which is attributed to the abundance of bile acid converting bacteria such as *P. hiranonis* in the canine gut.^25^ To isolate *bai*-containing bacteria with anti-*C. difficile* properties, we screened dog fecal samples in Taurocholate Moxalactam Norfloxacin Fructose medium (TMNF). TMNF is a modified version of *Clostridium difficile* Moxalactam Norfloxacin (CDMN) medium, a less commonly used medium for isolating *C. difficile* from fecal and other complex biological and environmental samples.^26, 27^ TMNF was used since an ideal anti-*C. difficile* bacterium should survive the same antibiotic pressure that commonly predisposes CDI in human patients, i.e. dysbiosis-inducing beta-lactams and fluoroquinolones.^28^ Thus, the isolated bacterium could serve as an adjunct live therapeutic along with dysbiosis-inducing antibiotics to prevent CDI. Therefore, screening donor fecal samples in a beta-lactam (Moxalactam) and fluoroquinolone (Norfloxacin) containing medium (TMNF) could provide a selective advantage for the candidate bacterial strains screened in this isolation process. In addition, the primary bile acid TCA in the medium could favor the isolation of bile acid metabolizing bacteria from donor samples. For the current study, further selection of presumptive colonies on TMNF agar was based on proline aminopeptidase activity, a quick disc-based test with a special emphasis on *P. hiranonis*. Proline aminopeptidase positive bacteria of interest include the *bai*-containing species-*P. hiranonis*, *C. bifermentans*, *C. sordellii* and the pathogen, *C. difficile.*^29^

The TMNF screening method effectively yielded five bacterial isolates from 50 samples tested known to have *bai*-operon (10% isolation rate), including two isolates of *C. bifermentans*, two isolates of *P. hiranonis*, and one isolate of *C. sordellii*. As expected, this screening method also yielded three *C. difficile* isolates (6% prevalence) (**Table 1**). In parallel, the canine fecal samples were screened in Cycloserine Cefoxitin Fructose Agar (CCFA), a widely used *C. difficile* isolation medium. The isolation rate for bai-operon containing bacteria was 4.77% (isolates of *C. sordelli* and *C. bifermentans,* and no *P. hiranonis*) with this isolation method, even after an extended screening of 270 dog fecal samples using this medium (**supplementary table 1**). However, recovery of *C. difficile* was higher (23%) in this method. Out of five isolates of *bai*-containing bacteria recovered from TMNF medium, the data from a unique *P. hiranonis* isolate (named *P. hiranonis BAI-17*) with notable anti-*C. difficile* biotherapeutic potential is presented in detail in this manuscript to describe the additional steps involved in our screening pipeline.

**Table 1.** Bacterial isolates recovered from 50 random canine fecal samples using TMNF screening method. The presence or absence of of *bai* and/or *bsh* genes known to be present in the available genomes or literature for each bacterial isolate is listed.

| Bacteria | Isolation rate (%) | *bai* operon | *bsh* gene |
| --- | --- | --- | --- |
| *Bifidobacterium pseudolongum* | 2 | No | Yes |
| *C. bifermentans* | 4 | Yes | Yes |
| *C. difficile* | 6 | No | No |
| *C. paraputrificum* | 4 | No | Yes |
| *C. sordelli* | 2 | Yes | No |
| *Eubacterium tenue* | 2 | No | Putative |
| *P. hiranonis* | 4 | Yes | Yes |
| *Parabacteroides distasonis* | 2 | No | Yes |
| *Terrisporobacter glycolicus* | 2 | No | No |
| *Clostridium sporogenes* | 2 | No | Yes |

Phenotypically, *P. hiranonis BAI-17* formed irregular translucent to rough grey colonies with serrated edges, very similar to *C. difficile* on blood agar plates. As previously mentioned, the proline aminopeptidase test is considered a rapid test for *C. difficile* does not distinguish *P. hiranonis* from *C. difficile.*^30^ The lack of fluorescence under UV light is found to be an easy test that differentiated *P. hiranonis BAI-17* from *C. difficile* (**Supplementary figure 1**).

**Figure 1.**
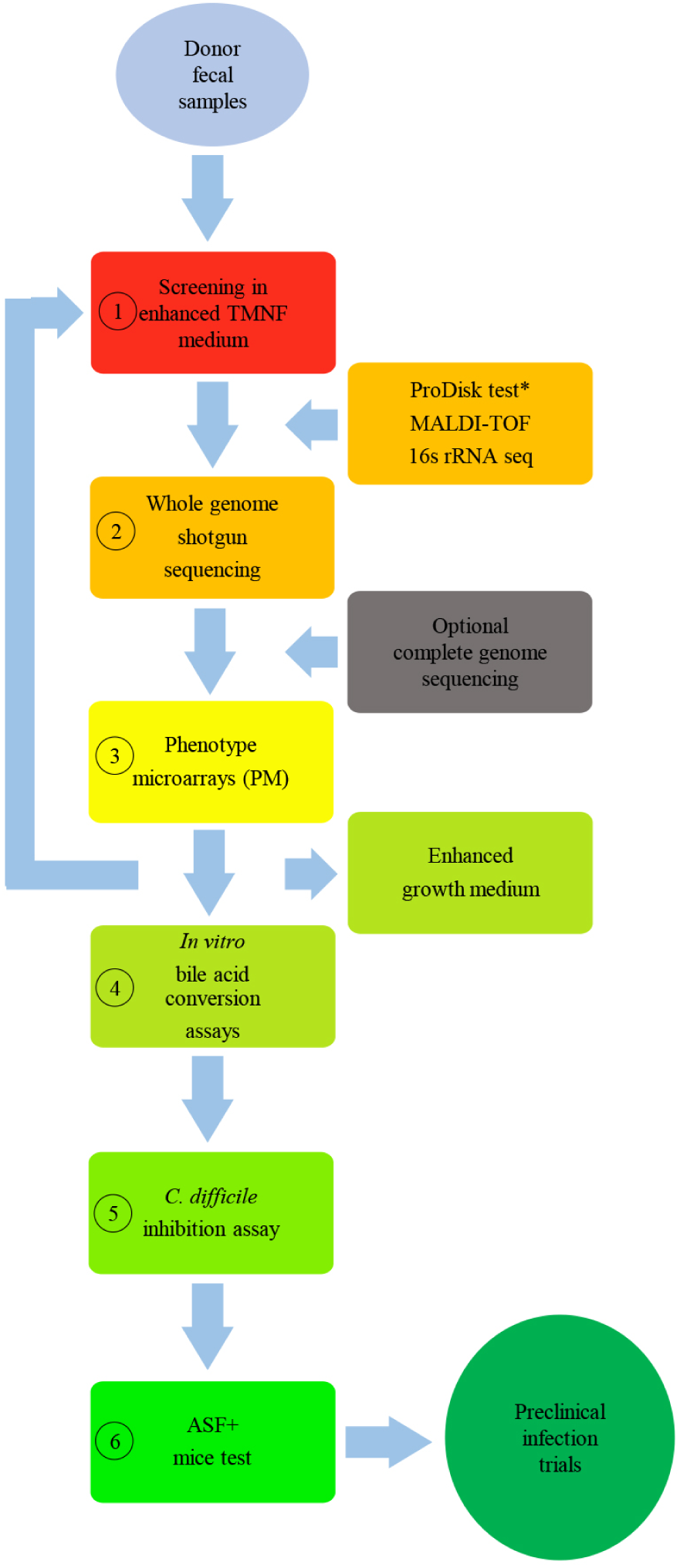
Overview of the screening pipeline for *bai*-containing bacteria with anti-*C. difficile* biotherapeutic potential. 1) screening in TMNF medium enables isolation of bile acid converting bacterium with similar growth condition as with *C. difficile*; 2) whole genome shotgun sequencing for identifying genes associated with bile acid tranformation, virulence, antibiotic resistance and gut colonization; 3) Phenotype Microarrays (PM) allows formulation of specific screening media for specialized screening of a particular bacterium, preparation of an enhanced growth medium and predict the competative substrate utlization of the candidate bacterium compared to *C. difficile*; 4) *in vitro* bile acid conjugation assay and 7-α dehydroxylation assay confrims and charecterize the bile acid tranformation ability of the candidate bacterium; 5) ASF^+^ mice test confirms *in vivo* bile acid tranformation and colonization ability of the candidate bacterium in the dysbiotic gut environment and assess the pathogenisity of the particular bacterium. * ProDisk test, if included screens specifically *P. hiranonis, C. sordelli,* and *C. bifermentans*

### *P. hiranonis BAI-17* possesses a unique combination of both *bsh* and *bai* genes with the absence of known toxin genes

An initial Illumina MiSeq-based whole-genome shotgun (WGS) sequencing of *P. hiranonis BAI-17* confirmed the presence of the *bai* operon and other bile acid metabolism-related genes. The annotation of the contigs by Rapid Annotation using Subsystem Technology (RAST) and mapping against *P. hiranonis* DSM 13275 in Geneious Prime software (Biomatters Inc, San Diego, CA) revealed both the presence of a unique *bai* operon involved in primary bile acid to secondary bile acid conversion (7-α dehydroxylation) and the *bsh* gene (choloylglycine hydrolase), which is responsible for the hydrolysis of conjugated bile acids required prior to 7-α dihydroxylation (**Figure 2**). This observation is remarkable because no other gut bacteria (especially Clostridiales) are known to have both critical genes of bile acid metabolism in a single genome.^31–33^ Further analysis of the currently available *P. hiranonis* genome sequences (one each of human and dog origin) in GenBank resulted in the same observation of *bai* and *bsh* presence in those strains.^29, 34^

**Figure 2.**
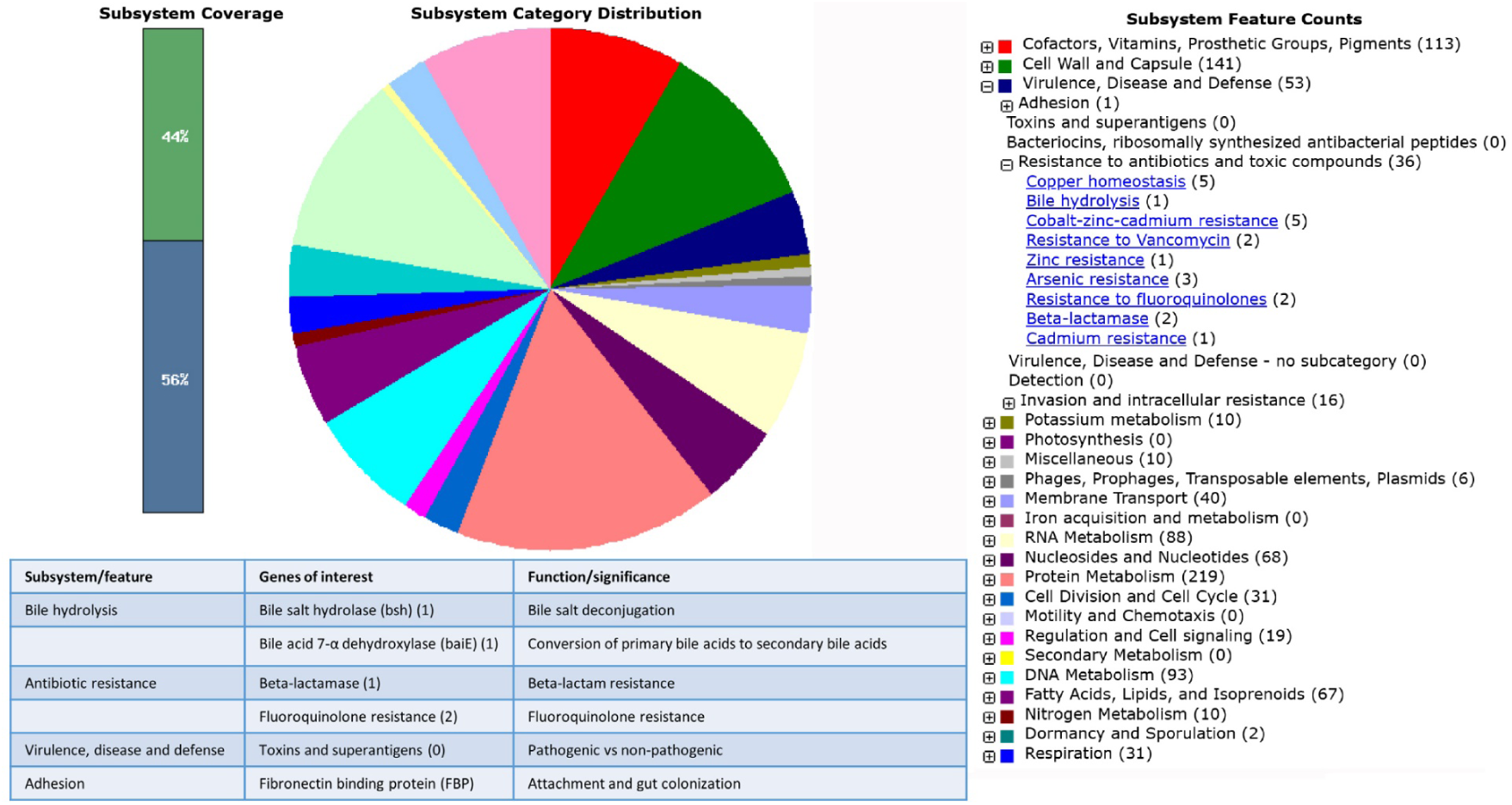
Whole-genome shotgun sequencing of *P. hiranonis BAI-17* reveals bile acid conversion genes and the absence of potential toxins and other virulence factors. The bacterial DNA isolated from a pure culture of *P. hiranonis BAI-17* was sequenced using the Illumina MiSeq platform. The assembled contigs were annotated using the RAST pipeline, and the genome and subsystems were visualized in Seed Viewer. The subsystem category distribution and feature counts are depicted and listed on the right and left of the panel, respectively. The table summarizes the major *P. hiranonis BAI-17* genes and functions that are favorable from an anti-C. difficile biotherapeutic perspective.

Having both *bsh* gene and *bai* operon in a single bacterium has immense therapeutic significance. Such bacteria could perform the functions of two different groups of metabolically dependant commensal bacteria in a dysbiotic gut of a CDI patient to restore normal bile acid metabolism. Thus, this bacterium alone could be supplemented as an adjunct prophylactic treatment during the administration of beta-lactams and fluoroquinolones in susceptible patients or could be administrated as a replacement or enrichment for FMT.

The WGS results also confirmed the presence of genes associated with beta-lactam and fluoroquinolone resistance in *P. hiranonis BAI-17* derived from the screening process (**Figure 3 and supplemental sequence data**). In addition, the intestinal colonization factor-fibronectin binding protein (FBP) was also identified within the *P. hiranonis BAI-17* genome. No known toxins or other virulence factors were identified, suggesting that this bacterium is presumptively non-pathogenic.

**Figure 3.**
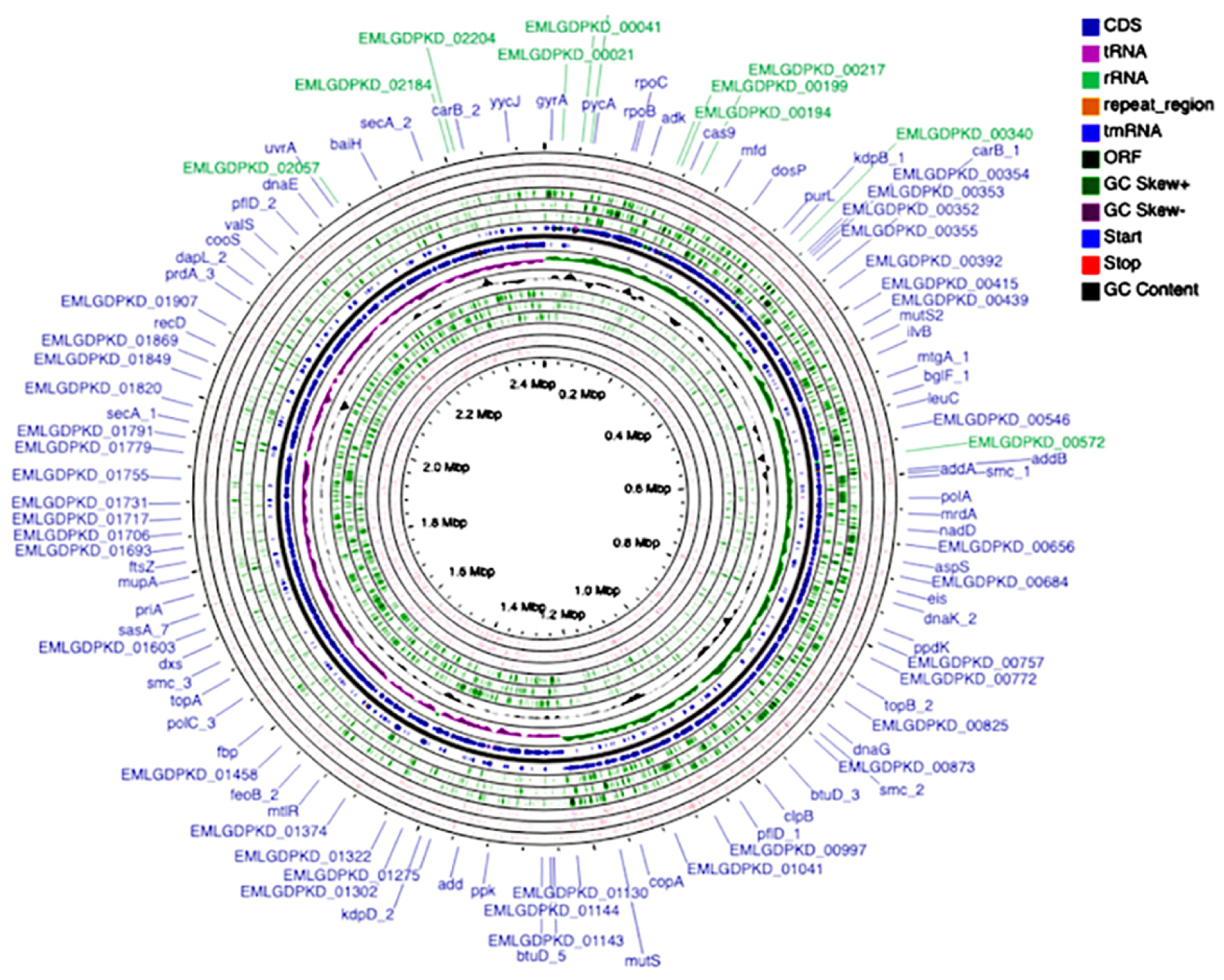
**Circular complete genome plot for *P. hiranonis BAI-17***: circular plot with tracks showing the CDS, tRNA, rRNA, repeat_region, tmRNA, ORF, GC Skew-, GC Skew+ , start codons, stop codons, and GC content.

### *P. hiranonis* is closely related to the bile acid converting gut bacterial species - *C. scindens* and *C. hylemonae*

The whole-genome shotgun sequencing step included in the screening method putatively examines the presence of *bai*, *bsh*, and other virulence and antibiotic resistance-related attributes in the candidate bacterium. However, as an optional step, a complete genome sequencing of *P. hiranonis BAI-17* was performed. The hybrid assembly using Illumina and Nanopore sequencing generated a single contig composed of 2,511,424 bp. The annotation of the genome using Prokka produced 2,307 total genes and 2,185 coding sequences. This is comparable to the only other publicly available *C. hiranonis* DSM 13275 genome composed of 2,521,899 bp, which has 2,363 genes and 2,239 CDS, respectively.^32^ A circular genome graph of *P. hiranonis BAI-17* with complete genome features is given in **Figure 3.**

### *Comparative Phylogenomic Analysis of* P. hiranonis BAI-17

The comparative phylogenomic analysis of *P. hiranonis BAI-17*, *P. hiranonis DSM 13275, C. scindens, and C. hylemonae strains* are showed in **Figure 4A.** As shown in **Figure 5A**, this analysis formed two major clades. *P. hiranonis BAI-17*, *P. hiranonis DSM 13275* and *C. hylemonae* strains clustered together, whereas *C. scindens ATCC35704* and *C. hylemonae DSM15053* were the closest species to *P. hiranonis BAI-17*.

**Figure 4.**
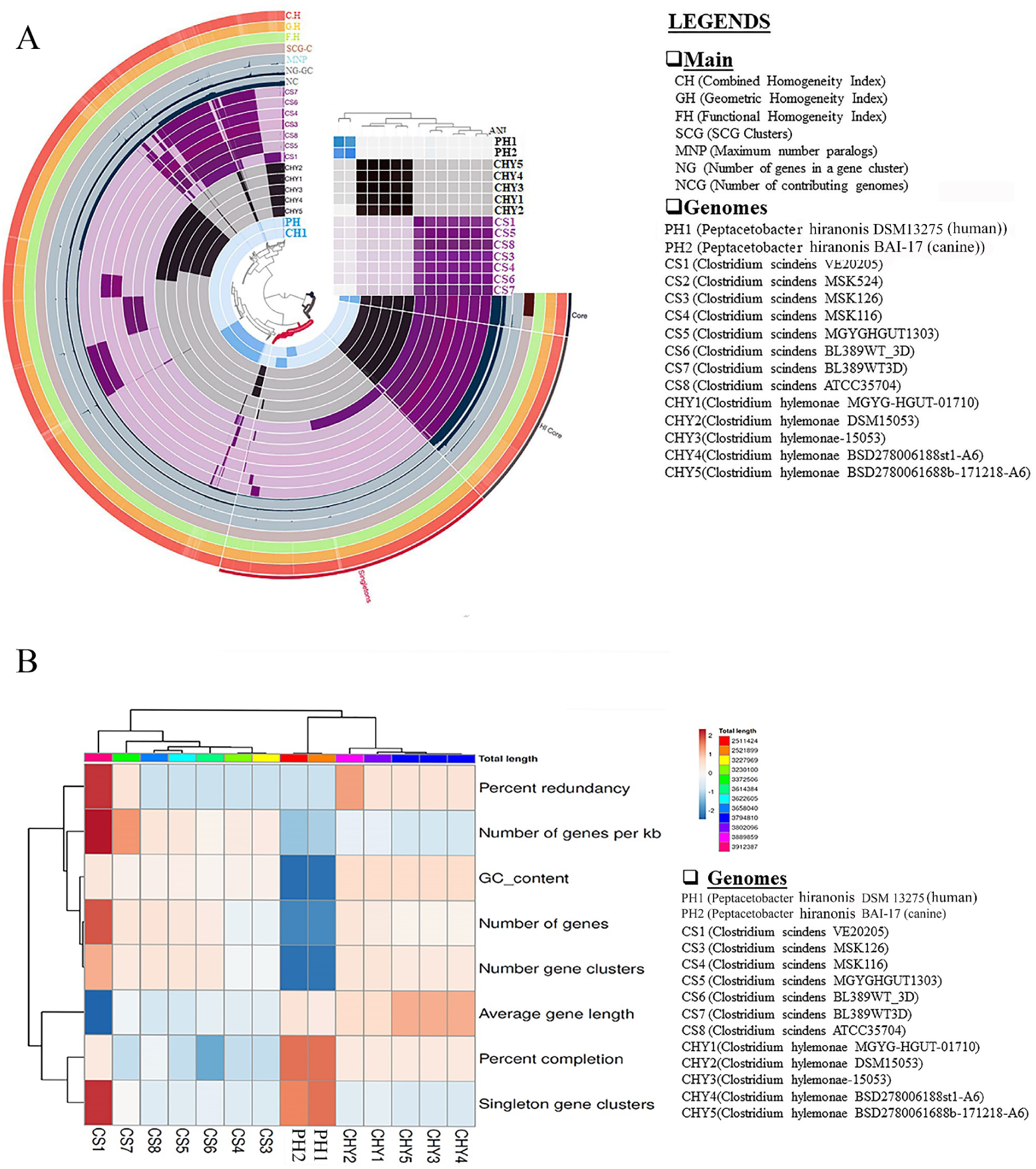
Comparative pangenome analysis of *P. hiranonis BAI-17*: The Pangenome analysis of *P. hiranonis BAI-17*, *P* (canine)*. hiranonis* DSM 13275 (human), seven *C. scindens* strains and four *C. hylemonae* strains (A) The genomic characteristics highlighting the number of singleton gene clusters, average gene length, number of gene clusters, number of genes, GC_ content, number of genes per kb, percentage redundancy and completion of each genome shown in a heatmap (B).

**Figure 5.**
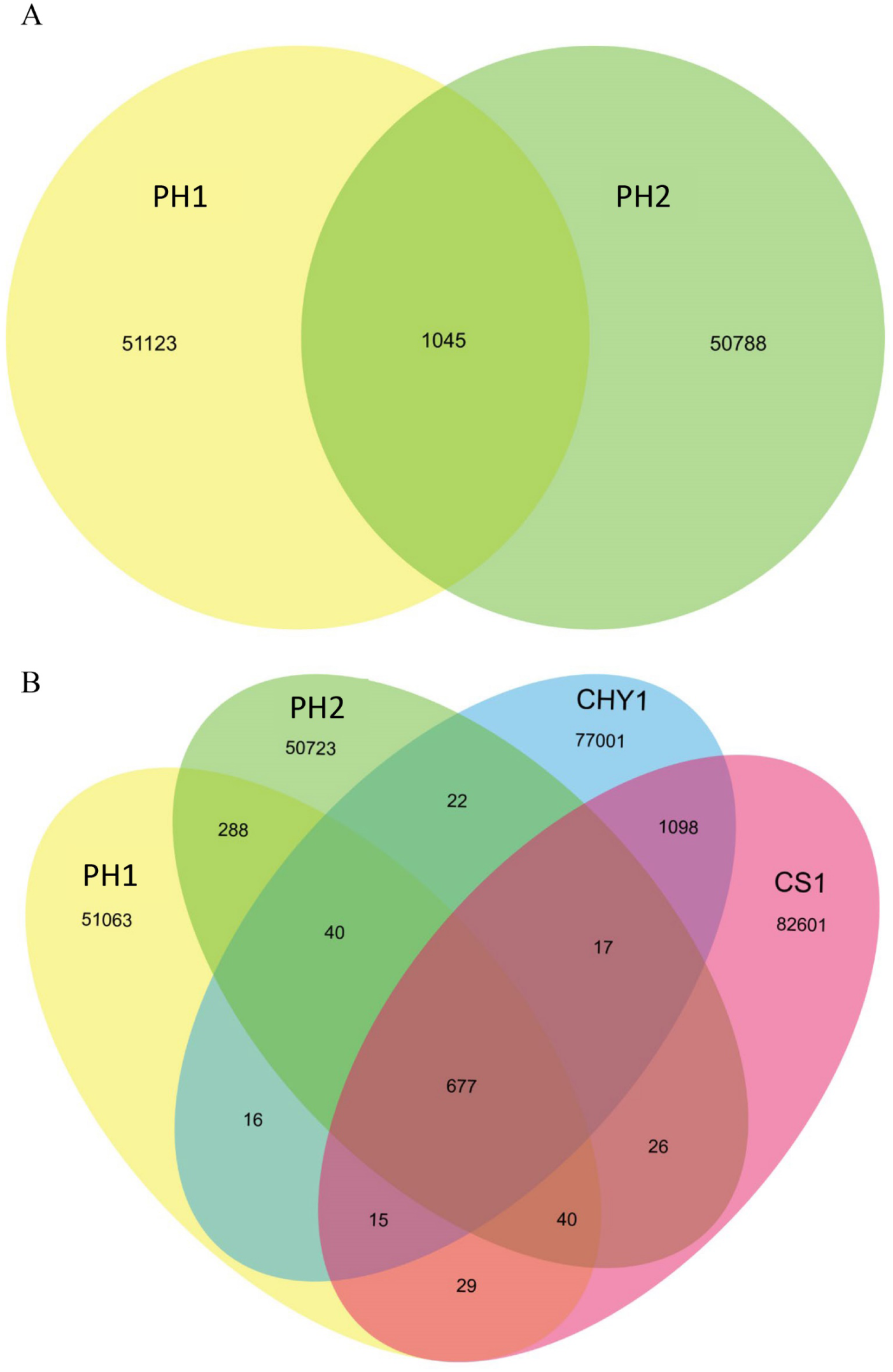
Comparative pangenome analysis of *P. hiranonis BAI-15*: **A)** Comparative analysis of gene families between canine origin *P. hiranonis BAI*17 and human origin *P. hiranonis* DSM 1327; **B)** Gene family analysis of *P. hiranonis BAI*17 (PH2), *P. hiranonis* DSM 13275 (PH1), *C. hylemonae* MGYG-HGUT-01710 (CHY1), and *C. scindens* VE20205 (CS1) showing the number of unique, shared and core genes within the genomes.

### Pangenome analysis

A pangenome analysis was performed using the following 14 bile degrading bacterial strains: *P. hiranonis BAI-17*, *P. hiranonis DSM 13275*, *Clostridium scindens BL389WT3D*, *Clostridium scindens VE202-05*, *Clostridium scindens MSK.1.26*, *Clostridium scindens BL-398-WT-3D*, *Clostridium scindens MGYG-HGUT-01303*, *Clostridium scindens MSK.1.16*, *Clostridium scindens ATCC35704, Clostridium hylemonae MGYG-HGUT-01710, Clostridium hylemonae DSM15053, Clostridum hylemonae DSM-15053, Clostridium hylemonae BSD2780061688st1_A6, and Clostridium hylemonae BSD2780061688b-171218_A6.* As expected, *P*. *hiranonis BAI-17* clustered with the publicly available human *P. hiranonis DSM 13275* and formed a clade with the *C. hylemonae* strains (**Figure 4A**). Also, all *C. scindens* strains clustered together, forming one clade. The attached bar plot (**Figure 4A**) shows that the average nucleotide identity (ANI) of the strains clustered according to the species-level variation. Comparatively, the lowest GC-content was recorded in *P. hiranonis* BAI-17 (0.3125) and *P. hiranonis DSM 13275* (0.3128) with the highest percentage completion (100) for both *P. hiranonis BAI-17* and *P. hiranonis* DSM 13275. In contrast, the highest GC-content was registered by the *C. hylemonae* strains (0.4895). There are approximately 2,205 genes observed in *P. hiranonis BAI-17* with an average gene length of 955.48. For comparison, the human *P. hiranonis DSM 13275* has approximately 2,246 genes with an average gene length of 941.50. Additionally, the average number of genes observed in *C. scindens* and *C. hylemonae* strains were 3673.86 and 3565.2, with an average gene length of 875.78 and 981.85, respectively. *P. hiranonis DSM 13275* has 2,122 gene clusters, which is higher than *P. hiranonis BAI17* having 2,105 gene clusters. However, the number of gene clusters for *P. hiranonis DSM 13275* was not higher than the average number of gene clusters in *C. scindens* (3401.26) and *C. hylemonae* (3416) strains. Out of this, 319 and 340 singleton gene clusters were observed in *C. hiranonis* BAI17 and *C. hiranonis DSM 13275* respectively. The lowest average number of singleton gene clusters were recorded in *C. scindens* (81.71) and *C. hylemonae* (10.6).

### Genome diversity within human and canine P. hiranonis strains

Since members of the genus *Peptacetobacter* have highly divergent genomes, the genome of canine *P. hiranonis BAI-17* was compared to the human *P. hiranonis DSM 13275* to determine the diversity between *P. hiranonis* species. This analysis demonstrated that approximately 50,788 genes or gene families were unique to *P. hiranonis BAI-17*, while 51,123 gene families were unique to *P. hiranonis* DSM 13275. Approximately 1045 gene families were shared between our *P. hiranonis BAI-17* and *P. hiranonis DSM 13275* (**Figure 5A**).

In addition, the genomes of *P. hiranonis BAI-17*, *P. hiranonis* DSM 13275*, C. hylemonae MGYG-HGUT-01710, and C. scindens VE20205* were compared. Results indicated that, 51,063 gene families were unique to *P. hiranonis* DSM13275, 50,723 gene families were unique to *P. hiranonis BAI-17*, 77,001 gene families were unique to *C. hylemonae MGYG-HGUT-01710 and* 82,601 gene families were unique to *C. scindens VE20205*. Approximately 677 core genes (gene families common to all four strains) (**Figure 5B**).

### Phenotype microarrays provide *P. hiranonis* growth requirements and substrate utilization profile of therapeutic importance

Phenotype microarrays (PM) are one of the most comprehensive microbial cell metabolic profiling techniques available.^35^ Biolog Phenotype Microarray metabolic panels (PMs 1–8) are composed of 200 assays of carbon source metabolism, 400 assays of nitrogen source metabolism, 100 assays of phosphorus and sulfur source metabolism, and 100 assays of other biosynthetic pathways.^35^ We performed PM was performed on *P. hiranonis BAI-17* to determine the nutrient utilization profile, and to to prepare an enhanced nutrient medium that can be used for large-scale culturing of this bacteria for screening, and preclinical and clinical studies. In addition, PM profiling of *P. hiranonis BAI-17* enabled comparison of the results to the published *C. difficile* PM profiles for shared metabolites of pathologic or therapeutic importance.^36^

The results indicate that 113 of 760 substrates in eight PM plates were utilized by *P. hiranonis BAI-17* (**Figure 6**). The most notable substrates that increased the growth of *P. hiranonis BAI-17* were glycyl-L-glutamic acid (64% increase in growth compared to negative control), pyroglutamic acid (54%), L-glutamine (52%), D, L, α-amino-caprylic acid (49%), octopamine (49%), 2-aminoethanol (ethanolamine) (47%), p-hydroxy phenylacetic acid (p-HPA) (47%), and a group of dipeptides (up to 55% increase in growth). Based on the PM analysis, a selected subset of these substrates (glycyl-L-glutamic Acid, L-glutamine, 2-aminoethanol, 2-amino caprylic acid, and p-hydroxy phenylacetic acid) was included in culture media for further screening and propagation of *P. hiranonis*.

**Figure 6.**
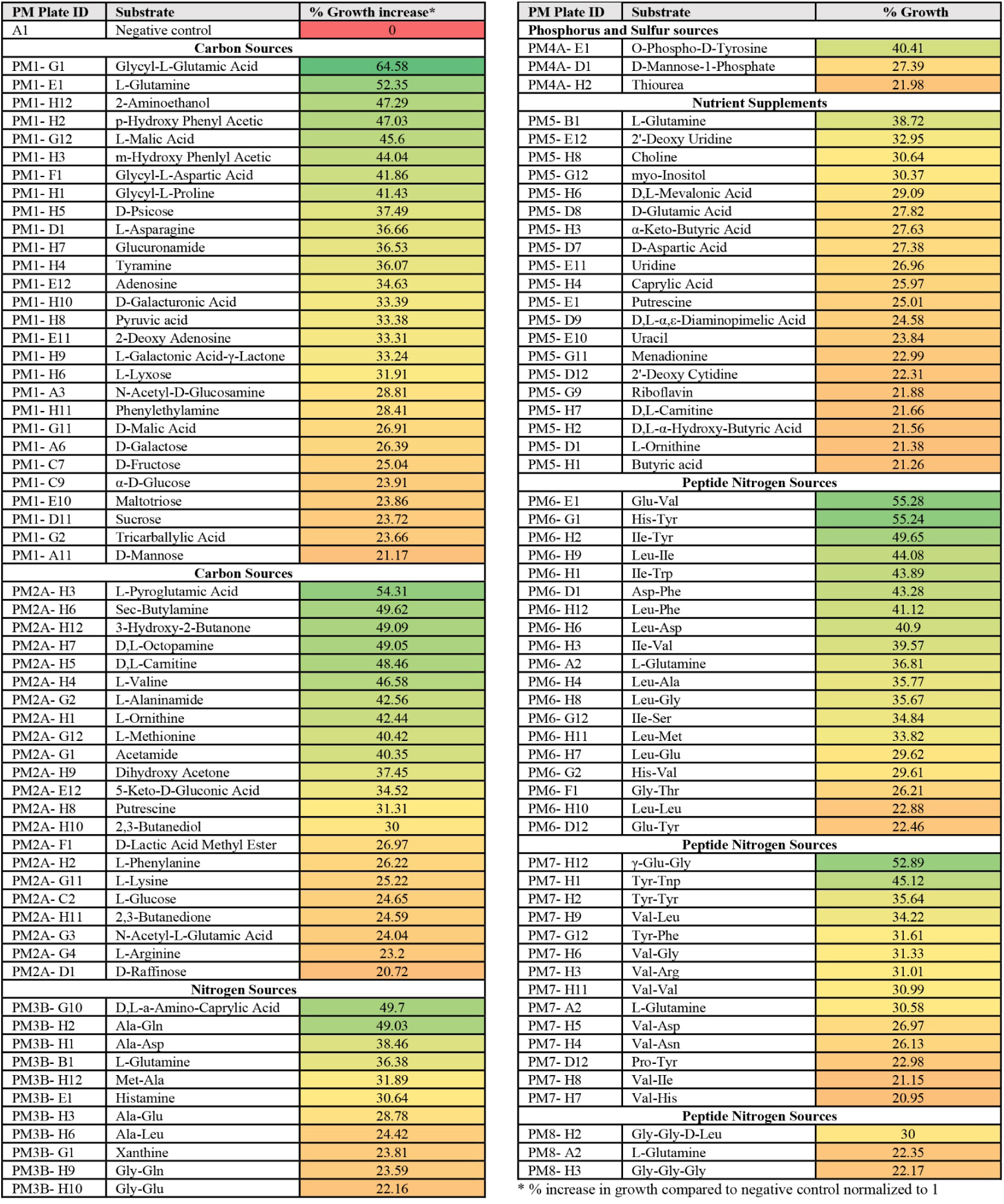
Substrate utlization of *P. hiranonis BAI-17* in Biolog Phenotype Microarrays: The nutritional phenotype of *P. hiranonis BAI-17* strain was determined using Biolog Phenotype MicroArray panels 1–8. Each row represents a nutritional phenotype that increased the bacterial growth by at least 20% of the negative control (i.e., bacterial growth equivalent to 120% of the negative control). The nutrient utilization for each panel is represented as a heat map in descending order of % increase in growth compared to the negative control (E.g., 169% of negative control means’ 69% increase in growth’).

The observation that *P. hiranonis BAI-17* utilizes ethanolamine is significant. The ability to utilize ethanolamine has been shown to provide a nutritional growth advantage over other enteric bacteria since ethanolamine is abundant in mammalian gut epithelial cell membranes.^37, 38^ Most importantly, ethanolamine catabolism has been demonstrated to induce the expression of virulence genes in *C. difficile*.^37, 38^ Thus, the competitive ethanolamine utilization by *P. hiranonis BAI-17* may impact *C. difficile* virulence and growth in favorably from a bio-therapeutic perspective.^36, 37^ Furthermore, the utilization of p-HPA by *P. hiranonis BAI-17* is notable in terms of *C. difficile* pathogenesis. p-HPA utilization is an important step in *C. difficile* tyrosine metabolism and induction of toxic p-cresol production, which has an important role in *C. difficile* overgrowth and CDI pathology in the gut.^39–41^ Thus, competitive utilization of p-HPA by *P. hiranonis BAI-17* may significant impact *C. difficile* virulence in the gut. In short, PM method provides critical information to formulate an optimum growth medium for large-scale culturing of the candidate bacterium and to understand the therapeutically relevant competitive substrate utilization profile from an anti-*C. difficile* biotherapeutic perspective.

### *P. hiranonis BAI-17* exhibits robust bile acid deconjugation and 7-α hydroxylation activities and inhibits *C. difficile* growth and toxin production *in vitro*

As described above, the genomic analysis of *P. hiranonis BAI-17* identified both *bai* and *bsh* genes and predicted both bile acid hydrolysing and 7-α hydroxylation ability. To confirm the functional *bai* and *bsh* expression of *P. hiranonis BAI-17 in vitro,* a primary to secondary bile acid conversion (7-α dehydroxylation) and bile acid hydrolysis assays were performed separately. The results demonstrated that *P. hiranonis BAI-17* deconjugated taurocholic acid (TCA) (a predominant conjugated primary bile acid in gut) to cholic acid (CA) in BHIS medium within 48 h as evident by the complete disappearance of taurine conjugate and appearance of CA in the media (**Figure 7A**). This observation confirms the presence of a functional *bsh* gene in *P. hiranonis BAI-17*. Thus, *P. hiranonis BAI-17* need not depend on other gut commensals for initial bile acid hydrolysis of conjugated primary bile acids secreted from the liver.

**Figure 7.**
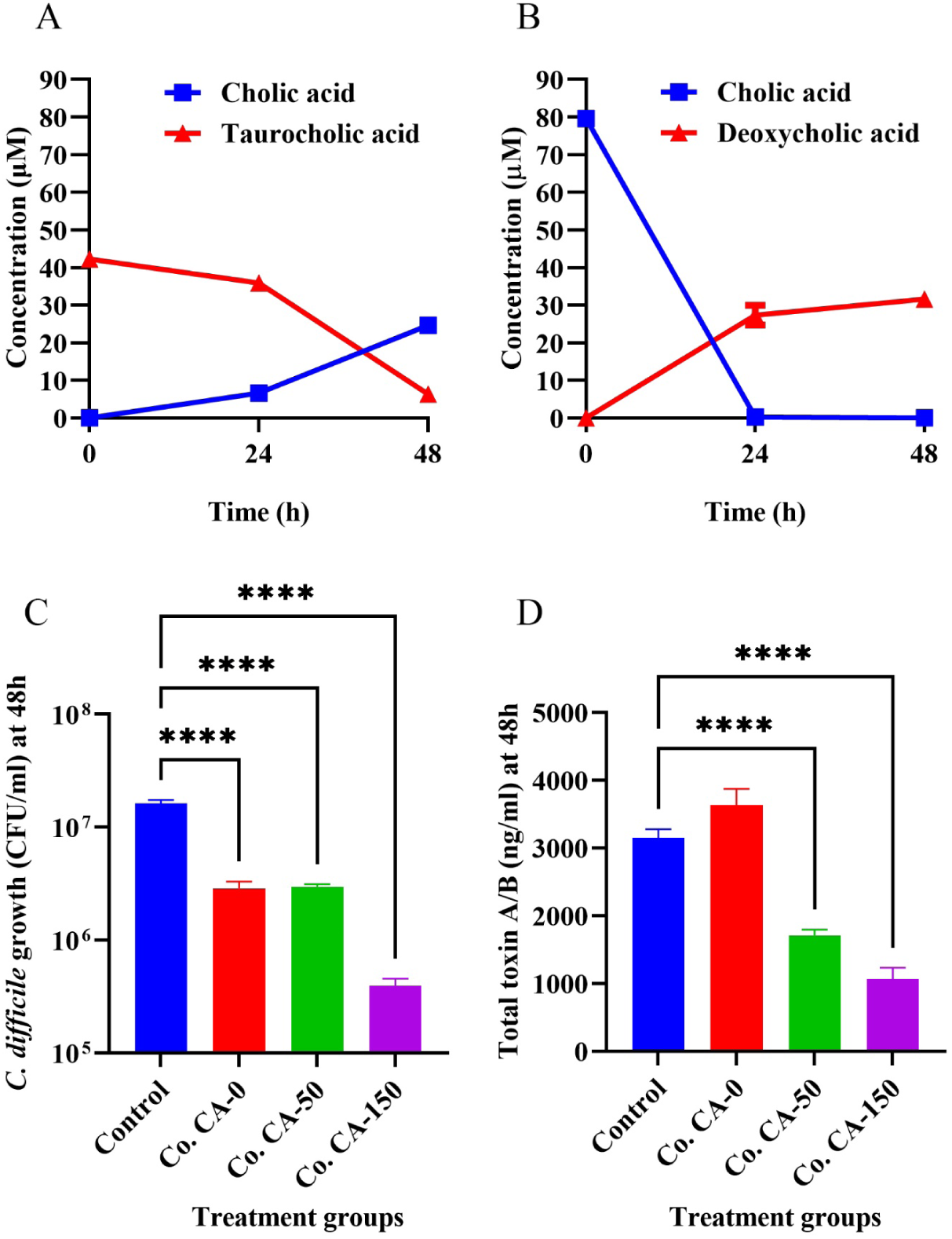
*P. hiranonis BAI-17* exhibits robust bile acid deconjugation (A) and 7-α dihydroxylation (B) ability and inhibits *C. difficile* growth *in vitro*: **A)** *P. hiranonis BAI-17* was cultured anaerobically in supplemented brain heart infusion (BHIS) broth with 50µM taurocholic acid, and the bile acids were quantitated from culture supernatants at 0, 24, and 48h using LC/MS technique. Taurocholate was completely deconjugated to cholic acid within 48h. **B)** *P. hiranonis BAI-17* was cultured anaerobically in supplemented brain heart infusion (BHIS) broth with 100µM cholic acid, and the bile acids were quantitated from culture supernatants at 0, 24, and 48h using LC/MS technique. Cholic acid was completely depleted to secondary bile acid DCA and other unmeasured metabolites by 24h; n=6 per experiment. **C)** *C. difficile* UK1 and *P hiranonis BAI-17* (10^5^ CFU/ml inoculum each) were cocultured anaerobically in BHIS broth with 0, 50, or 150µM cholic acid separately (n=6). *C. difficile* count at 48h was determined by anaerobic dilution and plating on blood agar supplemented with taurocholate. The Control group (blue bar) represents the growth of *C. difficile* alone. **D)** 48h culture supernatant from the coculture broth was assayed for total toxin A and B using quantitative ELISA. Groups: *C. difficile* alone (no coculture) (Control); *C. difficile* and *P. hiranonis* coculture with no i.e.. 0µM cholic acid (Co. CA-0); coculture with 50µM cholic acid (Co. CA-50); and coculture with 150µM cholic acid (Co. CA-150). ****p<0.0001

Further, an *in vitro* bile acid conversion assay was performed to confirm the functionality of the *bai* operon in *P. hiranonis BAI-17*. As expected, our *P. hiranonis* isolate converted cholic acid to deoxycholic acid within 24 h in BHIS medium as evident by complete disappearance of CA and appearance of its secondary bile acid (deoxycholic acid-DCA) (**Figure 7B**). The molar difference between the products and substrates in both deconjugation and 7-α hydroxylation assays indicate the formation of intermediate metabolites.^32, 42^ In summary, the *in vitro* bile acid assay together with comparative genomic analysis of available human and canine isolates establishes *P. hiranonis* as a bile acid deconjugating and 7-α hydroxylating bacterial species that has immense therapeutic potential.

Since the primary purpose of the screening pipeline is to identifying potential candidates for preclinical CDI infection trials, an *in vitro C. difficile* coculture experiment was included in the screening protocol. The *bai* operon-containing bacteria, *C. scindens* is a well-established candidate bacterium known to prevent *C. difficile* growth and virulence *in vitro* and *in vivo* in a bile acid-dependent manner.^19, 43, 44^ Although less widely isolated in laboratories, it is known that *P. hiranonis* is associated with notable colonization resistance of *C. difficile* in dogs.^25^ Since the bile acid metabolizing activity of *P. hiranonis BAI-17* has already been established from the previous steps described above, we expected to find similar results with *P. hiranonis in vitro*. Indeed, the *in vitro* coculture experiment demonstrated that *P. hiranonis BAI-17* inhibits *C. difficile* growth in the presence of micromolar concentrations of cholic acid (CA) in a dose-dependent manner (**Figure 7C**) (p<0.005). This could be attributed to production of the secondary bile acid (deoxycholic acid (DCA), in this assay) with or without the secretion of other antibacterial molecules inhibitory to *C. difficile* growth.^44, 45^ A concomitant reduction *C. difficile* toxins (total toxin A and B) was also observed in a cholic acid-dependent manner (p<0.005). Interestingly, in the absence of cholic acid, the toxin production was either unchanged or increased despite the reduction in bacterial growth, underscoring the bile acid dependent anti-*C. difficile* properties of *P. hiranonis BAI-17* (**Figure 7D**).

### ASF^+^ mice experiment confirms *in vivo* bile acid conversion, potential non-host specific gut colonization ability and safety of *P. hiranonis BAI-17*

For the final step of the screening pipeline, a mice experiment was designed to determine if the results obtained *in vitro* could also be observed *in vivo*. This experiment quantitatively confirms the ability of a bacterium to perform 7-α hydroxylation in the gut environment. Since, conventionally reared mice (Conv-R) harbor a complex microbiome, which may contain multiple bile acid metabolizing bacteria, testing a bacterium for *bai* activity in relation to the gut bile acid profile is challenging. At the same time, using germ-free or gnotobiotic mice also has several limitations, including the lack of a functional ’host gut-microbiome interface’, compromised immune system, diminished physiologic functions, and the need for costly animal care facilities. The interplay between the host*-*gut microbiome interface and the intestinal metabolites, specifically bile acids, is complex and requires a minimalistic but functional model for such studies.^16, 24^

The Altered Schaedler flora (ASF) is a murine bacterial consortium comprised of 8 known species ^46^. Unlike germ-free or mono-associated animals with deficiencies in immune system development and function, mice colonized with the ASF possess a stable microbiome and normal immune and physiologic functions compared to Conv-R.^46, 47^ Moreover, ASF mice can be maintained in a non-gnotobiotic environment for several weeks, since the defined resident microflora is stable.^46, 47^

To establish the suitability of ASF mice for assaying bile acid metabolism *in vivo*, the annotated genomes of all eight ASF consortium members^48^ are examined and confirmed that none of the ASF contains the *bai* operon. In contrast, four of the ASF members were predicted to encode *bsh* gene. These observations indicate that the ASF is a suitable model to quantify and assess 7-α dehydroxylation ability by heterologous microorganisms *in vivo*. Also, these experiments could also provide *in vivo* safety and toxicity assessment of the candidate bacterium, confirming the genome-wide virulence screening performed in the preceding steps of the screening pipeline. The same experiments could also provide insight into the inter-host colonizing ability (canine strain in this case) of the candidate bacterium that has significance in treating CDI in different mammalian species.

### P. hiranonis BAI-17 *readily colonizes the ASF mouse gut*

Results showed that a single oral gavage of 10^5^ CFU of *P. hiranonis BAI-17* in adult ASF mice resulted in functional gut colonization of this bacterium. This observation is based on detection and visualization of *P. hiranonis BAI-17* DNA and RNA in the gut 3-weeks post-inoculation using qRT-PCR and RNAscope *in situ* hybridization methods, respectively. Positive amplification was observed in the colonic contents of all ASF mice inoculated with *P. hiranonis BAI-17* (ASF-PH mice) euthanized at 9-weeks of age (**Figure 8D**), while no *P. hiranonis BAI-17* specific amplification was observed in control ASF mice. In addition, RNAscope *in situ* hybridization of colonic tissue sections from mice euthanized 3- weeks post-inoculation revealed localization of *P. hiranonis BAI-17* in both luminal contents and within the tight mucus layer of the colonic mucosa (**Figure 8A, B &C**). No positive signal was observed in control ASF mice.

**Figure 8.**
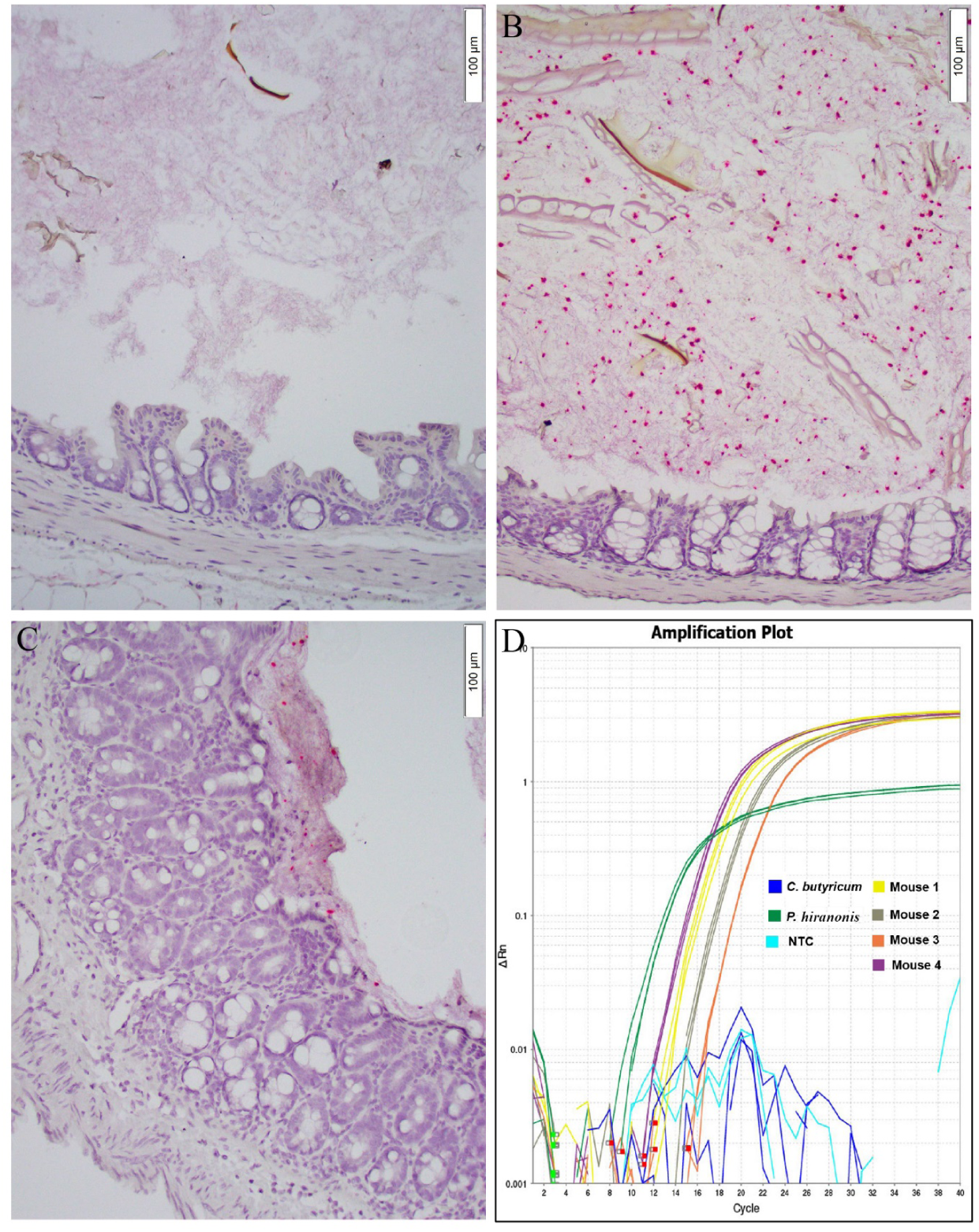
*P. hiranonis BAI-17* colonizes ASF mice: Six-week-old ASF mice were inoculated via single oral gavage with 250 µl of 10^5^ CFU/mL *P. hiranonis BAI-17* or PBS. The animals were euthanized, and the large intestine with contents were collected after 3-week post-inoculation. *P. hiranonis BAI-17* bacteria (red signal) was demonstrated by RNAscope *in situ* hybridization technique using custom-made specific RNAScope probes. (**A):** cecum-negative control (ASF); **B)**: cecal luminal contents of an ASF mouse colonized with *P. hiranonis BAI-17*; **C):** depiction of tight mucus in the colon an ASF mouse colonized with *P. hiranonis BAI-17* **D)** Colonization with *P. hiranonis BAI-17* was initially confirmed by TaqMan based qPCR using fresh fecal pellets collected from individual mice three weeks post-inoculation (result from four ASF shown). DNA from *P. hiranonis BAI-17* and *Clostridium butyricum,* along with a non-template control (NTC) were included as test controls.

### P. hiranonis BAI-17 *colonization does not produce adverse effects in mice*

Clinical, gross, and histologic examination revealed no adverse enteropathologic changes in ASF mice colonized with *P. hiranonis*. Histopathology did not reveal significant inflammatory, toxic or degenerative changes in the gastrointestinal system of ASF and ASF-PH mice (data not shown).

### P. hiranonis colonization alters the gut bile acid profile

As expected, the secondary bile acids DCA and LCA were not detected in ASF control mice, whereas a significant amount of these secondary bile acids was present in the colonic contents of ASF-PH mice (**Figure 9).** The DCA concentration in ASF-PH mice gut attained the same levels of CA indicating robust 7-α dihydroxylation activity of *P. hiranonis BAI-17*. This finding confirmed the *in vivo* bile acid converting ability of *P. hiranonis.* Most of the detected bile acids were unconjugated, indicating effective bile salt hydrolysis or ileal absorption of the remaining conjugated bile acids. Primary bile acids CDCA and α-epimer of MCA were significantly elevated in ASF mice compared to ASF-PH mice, presumably due to lack of 7-α dihydroxylation and conversion to secondary bile acids (p<0.05). Minimal CDCA with a high LCA concentrations in ASF-PH mice indicates a robust CDCA to LCA conversion. Thus, this study effectively demonstrated the *in vivo* bile acid conversion of our candidate bacterium. As detailed previously, the depletion of *bai*-containing bacteria (e.g., post-antibiotic treatment) is a major cause for bile-acid dysmetabolism and gut *C. difficile* overgrowth in susceptible individuals. Moreover, the ASF gut microbiota recapitulates a *bai*-depleted gut environment due to its limited microbiota composition. Thus, effective, non-pathogenic and functional colonization of *P. hiranonis BAI-17* by filling this depleted *bai* niche in the ASF mice validates the therapeutic potential of this bacterium. Therefore, follow-up preclinical and clinical trials using *P. hiranonis BAI-17* are likely to provide promising results as an effective anti-*C. difficile* biotherapeutic agent.

**Figure 9.**
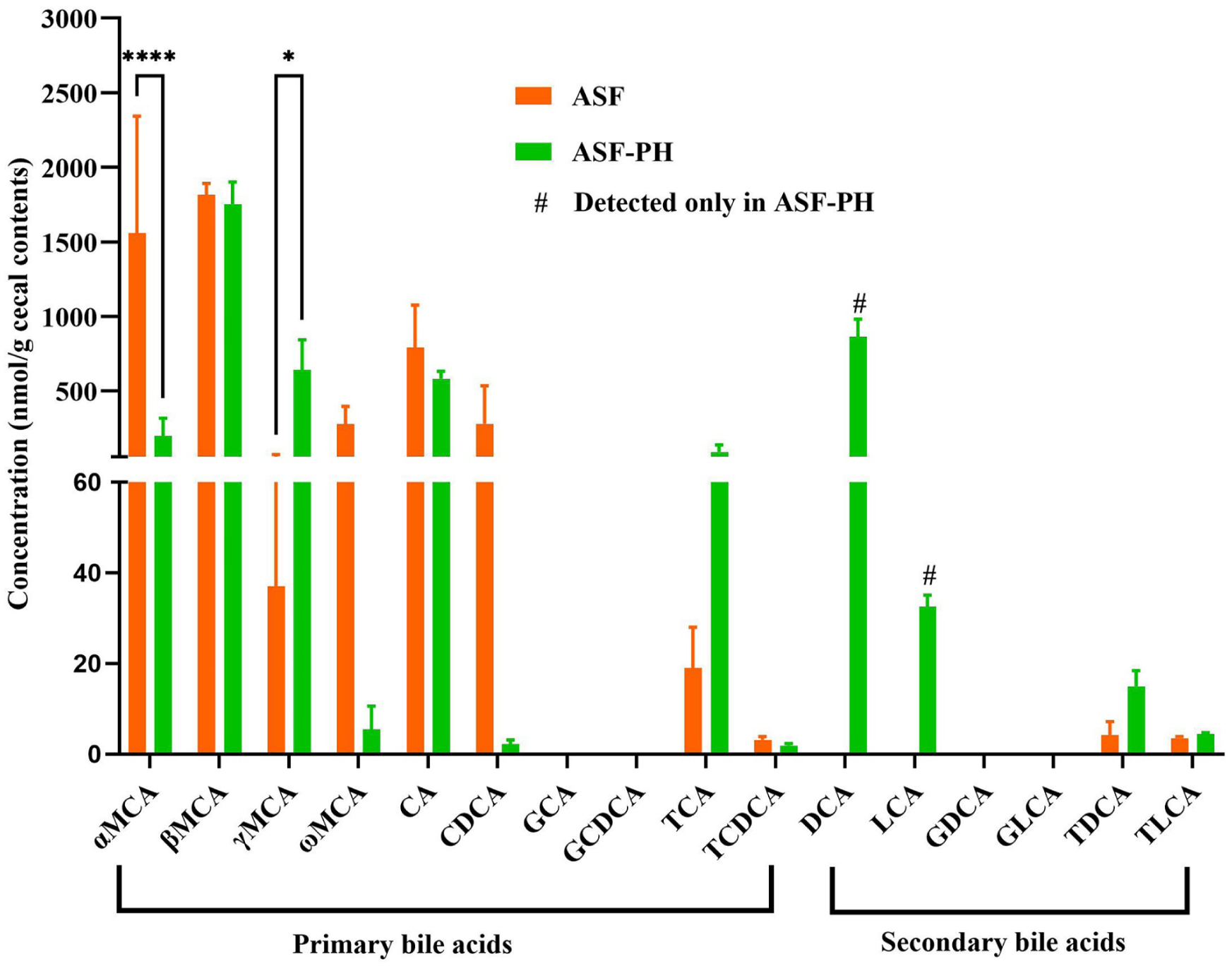
ASF^+^ test demonstrates the formation of secondary bile acids in the cecal contents from ASF mice colonized with *P. hiranonis BAI-17* (ASF-PH) but not in ASF control mice: Six-week-old ASF mice (n=6) were inoculated with single oral gavage with 250 µl of 10^5^ *P. hiranonis BAI-17* to generate ASF-PH mice. Cecal contents from both ASF-PH mice and age-matched control ASF mice were quantified for primary and secondary bile acids using targeted metabolomics analysis. Primary bile acids: MCA: muricholic acid; CA: cholic acid; CDCA: chenodeoxyxholic acid; GCA: glycocholic acid; GCDCA: glycochedeoxyxholic acid; TCA: taurococholic acid; TCDCA: taurochedeoxyxholic acid. Secondary bile acids: DCA deoxycholic acid; LCA; lithocholic acid; GDCA glycodeoxycholic acid; GLCA; glycolithocholic acid; TDCA taurodeoxycholic acid; TLCA; taurolithocholic acid. **#** detected only in ASF-PH but not in ASF control mice; ** p <0.005; **** p<0.0001.

## Discussion

Only a few species of 7-α dehydroxylating bacteria are known to be present in the mammalian gut. Almost all of them are members of the order Clostridiales and include *C. scindens*, *C. hylemonae*, *P. hiranonis*, some strains of *C. bifermentans,* and *C. sordelli*, and a few species of *Lachnospiraceae* and *Ruminococcaceae* families. These bacteria contain the unique *bai* operon in their genome that consists of the *baiABCDEF* gene cluster, which performs sequential steps in the colonic conversion of primary bile acids to secondary bile acids in mammals, including humans.^17^ Almost all the 7-α dehydroxylating bacteria are fastidious slow-growing anaerobes with poorly known growth characteristics and virtually non-existent specific isolation and culture protocols. Among these bacteria, *C. scindens* is the only one that has been characterized to a greater extent and been used for current research on bile acid metabolism and CDI precision microbiome therapy.^20, 43, 49, 50^ Hence, identifying new culturable strains of *bai*-containing bacteria with robust bile acid converting ability is critical from a biomedical research and therapeutic perspective. For achieving this goal, the development of a more defined and utility-specific screening pipeline is required. Therefore, a pipeline was developed in this study that specifically screens *bai-*containing bacteria that have a defined anti-*C. difficile* therapeutic potential.

The screening pipeline detailed in this study is specifically intended for isolating live biotherapeutic bacterial candidates for precision microbiome-based CDI prophylaxis and treatment. To test this fecal screening pipeline, dogs were used as model donors, since they are inherently resistant to clinical CDI despite asymptomatic *C. difficile* carriage.^25^ In addition, dogs are known to carry a high proportion of bile acid converting bacteria specifically *P. hiranonis* in their gut.^51^ Moreover, *P. hiranonis* colonization is inversely associated with gut dysbiosis and *C. difficile* carriage, a parameter clinically used for assessing the gut dysbiosis index in dogs.^25, 52, 53^ However, this pipeline described in this study has the flexibility to use any donor feces for isolating *P. hiranonis* and other *bai*-containing bacteria from any species.

Susceptible patients, especially the elderly, acquire CDI as a consequence of intensive antibiotic treatment, mainly beta-lactams and fluoroquinolones, administered for infections or surgical procedures. These antibiotics deplete the bile acid metabolizing bacterial flora in the gut, resulting in *C. difficile* gut colonization and overgrowth. In principle, CDI can be prevented by supplementing with robust bile acid metabolizing bacteria during administration of these antibiotics. Therefore, a *bai*-containing bacterium that can also withstand beta-lactam and fluoroquinolone pressure is a desirable target. Similarly, from a therapeutic viewpoint, a *bai*-containing bacterium with similar growth and nutrient requirements as *C. difficile* could have a competitive advantage when used alone or as enrichment for FMT. Thus, a medium containing a beta-lactam antibiotic (moxalactam), a fluoroquinolone (norfloxacin), and a conjugated primary bile acid (taurocholate), along with other nutrients (e.g., cysteine and blood), were selected for screening the *bai* containing bacteria from fecal samples. This method effectively yielded five bile acid-converting bacterial isolates of three different species from 50 fecal samples tested (10%), including one (*P. hiranonis BAI-17*) that passed all the subsequent screening steps involved in this isolation pipeline, making it into the infection trial phase. In addition, a widely used medium for isolating *C. difficile* from fecal samples (CCFA medium supplemented with taurocholate) was also investigated for isolating *bai*-containing bacteria. The *C. difficile* yield from this method was higher compared to the TMNF medium; however, the isolation rate of *bai*-containing bacteria was very low, i.e., 4.77% with no *P. hiranonis* isolates (**supplementary table 1**). Another caveat with this method is that the CCFA medium does not contain a fluoroquinolone antibiotic, and hence the candidate bacterium may not be suitable for prophylactic use along with fluoroquinolones.

The next step involved in the screening process was to verify genes in the candidate bacterium and their functional activity associated with functional bile acid metabolism, virulence, and antibiotic resistance. Screening for specific genes associated with therapeutic significance of several candidate probiotic bacteria has been conventionally done using detections via PCR amplification.^54^ However, our approach was to utilize next-generation sequencing, since it is now affordable and could provide comprehensive information on hundreds of genes and functional pathways simultaneously. Thus, we adopted WGS sequencing using the Illumina MiSeq platform for this purpose. Annotation of the assembled sequences confirmed the presence of *bai* genes in *P. hiranonis BAI-17*. However, identifying a *bsh* gene at this step further demonstrated the therapeutic value of the *P. hiranonis BAI-17* isolate. Generally, bile acid deconjugation in the gut is carried out by a set of bacteria that are primarily members of the phyla Bacteroidetes and Firmicutes. Notable bile salt hydrolysers in the gut include several species of *Bacteroides*, *Parabacteroides, Listeria, Bifidobacterium, and Lactobacillus.*^17^ Presence of a functional *bsh* has not been demonstrated yet in any other *bai* containing bacterial species. A few clostridial species, such as *C. perfringens* and *C. botulinum* contain *bsh* genes; however, they do not possess the *bai* operon.^31, 55^ The *bai* operon of *P. hiranonis* is comparable to that of other widely known bile acid converting bacteria namely *C. scindens* and *C. hylemonae,* which does not contain the *bsh* gene.^56^ The complete genome assembly of *P. hiranonis BAI-17* and comparative genome analysis data suggest a close genetic relationship between *P. hiranonis* (both canine and human isolates) and the well-known *bai*-containing bacteria C. *hylemonae,* and *C. scindens* although neither of them contains *bsh.* The functional activity of both *bsh* and *bai* of *P. hiranonis* is confirmed by the *in vitro* bile acid conversion tests, which includes a bile salt conjugation assay (for *bsh* activity) using the conjugated primary bile acid TCA and 7-α dehydroxylation assay (for *bai* activity) with the primary bile acid, CA. Further, the animal experiment (ASF^+^ mouse test) confirmed *in vivo* formation of secondary bile acid in *P. hiranonis* colonized mice suggesting that this bacterium could exert a therapeutic effect in the gut environment. Therefore, the novel ASF^+^ mouse model for specifically and quantitatively assessing bacterial 7-α dehydroxylation has been shown to be effective in meeting this purpose.

The WGS step included in the screening pipeline also provides a comprehensive picture of potential and putative virulence factors in the candidate bacterium. Members of the family Clostridiaceae, to which *P. hiranonis* belongs, are known for different exotoxins, including pore-forming toxins, phospholipase C, collagenase, large glucosylating toxins, binary toxins, and neurotoxins.^57^ The annotation of *P. hiranonis BAI-17* sequence assembly did not identify any known toxins or superantigens in its genome, which predicted the safety of this bacterium for therapeutic use. This observation was further confirmed *in vivo* by the absence of tissue lesions and adverse signs in ASF-PH mice post-inoculation. In addition, WGS sequencing also confirmed fluoroquinolone and beta-lactam resistance in the candidate bacterium, which corroborates with the selection pressure and screening process using the TMNF media. Since *C. difficile* is inherently resistant to these antibiotics, horizontal transfer of the resistance genes from the candidate bacterium to the pathogen is not a concern in this case.

WGS results also identified fibronectin-binding proteins (FBPs) in the *P. hiranonis BAI-17* genome. Fibronectin is widely expressed in the intestinal epithelium of mammals and is widely utilized by gut bacteria for mucosal adhesion.^58, 59^ FBPs in bacteria, including probiotic species such as *L. plantarum* and pathogens *C. difficile,* are involved in gut colonization and long-term retention of the bacterium in the intestinal lumen.^54, 58, 60–62^ Moreover, detection of FBP gene within the bacterial genome has been utilized to predict putative intestinal colonization of probiotic bacteria.^54^ Thus, although speculative, having this gene expressed could help *P. hiranonis BAI-17* to colonize the recipient animals independent of the host species. Interestingly, the *in vivo* experiment in this study demonstrated that a single oral dose of *P. hiranonis BAI-17* resulted in stable functional gut colonization of this bacterium in the adult ASF mice gut, which was confirmed by detection of this bacterium in the feces and production of secondary bile acids three weeks post-inoculation. This finding indicates the colonization potential of *P. hiranonis BAI-17* in a functionally and structurally dysbiotic gut (found ASF mice) of other non-hosts species including humans, horses and swine with CDI. Moreover, fibronectin binding using Fbp68 is crucial for *C. difficile* gut colonization.^61, 63–65^ Phylogenetic analysis of *P. hiranonis* FBP (including canine *P. hiranonis BAI-17* and DGF055142, and human DSM13275) using BLASTn revealed that the closest bacterial gene predicted was the FBP of *Romboutsia sp* (query cover 99%, E value: 0, and percentage identity 71%) and *C. difficile* (query cover 99%, E value: 0, and percentage identity 69%). Thus, it is also likely that *P. hiranonis BAI-17* would inhibit *C. difficile* colonization by competitive exclusion, adding to the therapeutic potential of this bacterium, and a question of interest in future studies.

Additionally, Biolog PM provided a set of metabolites that when supplemented in media enhances the *in vitro* growth of *P. hiranonis BAI-17*. This information enabled the formulation of an enhanced growth media for this bacterium. Although not tested in this study, it is also likely that the TMNF medium supplemented with these metabolites will allow an increased isolation rate of *P. hiranonis* strains. In addition, as detailed in results, utilization of ethanolamine, which is associated with virulent strains of *C. difficile,* could add to the biotherapeutic potential of *P. hiranonis BAI-17* as it may compete with *C. difficile* for this metabolite.^37, 38^ Phenotypic microarrays also indicated the utilization of pHPA by *P. hiranonis BAI-17*. pHPA is important in *C. difficile* pathogenesis as this pathogen converts pHPA to p-cresol. This toxic compound inhibits the growth of other bacteria in the gut and contributes to dysbiosis and *C. difficile* overgrowth.^66, 67^ *C. difficile* contains the *hpdBCA* operon involved in production of p-cresol from its precursor molecule pHPA.^39^ Interestingly, the WGS data revealed an absence of the *hpd* genes in *P. hiranonis BAI-17*, suggesting that pHPA utilization by this bacterium is not associated with p-cresol production. Thus, competitive utilization of pHPA by *P. hiranonis BAI-17* may result in fewer precursors available for p-cresol production, which will also contribute to its therapeutic significance.

In summary, the isolation pipeline, which involve fecal screening in TMNF media, WGS analysis, phenotype microarrays, *in vitro* bile conversion analysis, and *C. difficile* inhibition assays, as well as proof of concept *in vivo* studies (ASF^+^ mouse test) has been described. This protocol has been shown to efficiently isolate *bai*-containing bacteria from fecal samples, identify broad CDI-specific therapeutic characteristics, and determine the pathogenicity and safety. Thus, a bacterium that met all the criteria set can be passed to more extensive preclinical infection trials. In addition, the salient confirmation that *P. hiranonis* harbors both *bai* and *bsh* genes essential for bile acid conversion makes this bacterium therapeutically appealing. Altogether, the screening pipeline developed for bile acid converting bacteria revealed the unique anti-*C. difficile* biotherapeutic potential of *P. hiranonis* and these findings can be validated by follow-up infection studies in mice and subsequent clinical trials.

## Materials and methods

### Overview of the screening pipeline for *bai-*containing bacteria

The screening pipeline for isolating *bai*-containing bacteria with anti-*C. difficile* therapeutic potential has the following objectives: a) isolation of a *bai*-containing bacterium that can grow along with *C. difficile* and has similar competitive growth requirements and identical resistance pattern to bile acids, antibiotics and other gut derived chemicals as exhibited by *C. difficile*; b) genotypic screening of the candidate bacteria for virulence factors and other genetic attributes that has putative therapeutic advantage against *C. difficile*; c) phenotypic screening for metabolites that influence the growth of the candidate bacteria for identifying specific growth medium and future species specific screening medium; d) *in vitro* screening of the candidate bacteria for bile acid transformation activity and bile-acid dependant *C. difficile* inhibitory effect; e) *in vivo* screening for quantitative bile acid conversion ability in the gut environment in a *bai-* depleted animal model (ASF mice) which enables parallel assessment of gut colonization ability and potential tissue-organ toxicity. The overview of this pipeline and steps involved for meeting these objectives are depicted in **Figure 1**.

### Donor animals, sampling, and isolation of *bai-*containing bacteria from canine fecal samples

Random canine fecal samples were collected from multiple sources at Iowa State University (ISU) College of Veterinary Medicine: 1) small animal teaching hospital, Lloyd Veterinary Medical Center, 2) Department of Veterinary Pathology, and 3) ISU Veterinary Diagnostic Laboratory (VDL). Fifty random samples were used for screening in TMNF medium (modified *C. difficile* Moxalactam Norfloxacin medium with 0.1% taurocholate), and 250 samples were used for screening in CCFA medium as previously.^27, 30^ Approximately 1g of feces was resuspended in 10 mL of pre-reduced TMNF broth, mixed thoroughly and incubated in an anaerobic workstation (AS-580, Anaerobe Systems, Morgan Hill, CA, USA) for 7 days in anaerobic condition (0% oxygen, 5% hydrogen, 5% CO2 and 90% nitrogen at 37°C). After enrichment, the suspension was then centrifuged at 5000 rpm for 15 minutes. The supernatant was removed, the pellet was transferred to the anaerobic workstation, resuspended in 0.2 mL of TMNF broth, mixed thoroughly, and immediately plated onto pre-reduced nonselective blood agar (R01202, Fisher Scientific, Waltham, MA) and BHIS agar (Brain Heart Infusion supplemented with 0.5% yeast extract), and incubated at 37°C for 48 h in anaerobic condition as described above. After visible growth, colonies were tested for an L-proline aminopeptidase activity with PRO Disk (ThermoFisher Scientific, Waltham, MA, USA), to screen for *C. difficile* and bile acid converting bacteria *C. sordellii*, C. *bifermentans,* and *P. hiranonis*^29^) per the manufacturer’s instructions. Test positive colonies were subcultured onto CDMN agar and incubated for 24-48 h to obtain pure culture of each colony. After 48 h, the pure growths were harvested and subcultured in BHI broth for further experiments

### Bacterial identification

The identity of the candidate bacterium was determined using matrix-assisted laser desorption–ionization time-of-flight mass spectrometry (MALDI-TOF) and further confirmed by 16s rRNA PCR sequencing. The sample preparation and analysis were carried out as previously described.^68^ For PCR and sequencing, the broth culture was centrifuged at 5,000 x g at 4°C for 15 minutes. The supernatant was discarded, and the DNA was extracted from the bacterial pellet using the QIAamp BiOstic Bacteremia DNA Kit (Qiagen, Germantown, MD, USA). Extracted DNA was amplified by traditional PCR using universal 16S rRNA gene targeting primers. PCR was performed using Promega GoTaq® Master Mixes with the following thermocycler conditions: initial denaturation at 95°C for 3 minutes followed by 30 cycles of denaturation at 95°C for 30 seconds, annealing at 55°C for 30 seconds and extension at 65°C for 45 seconds and a final extension at 72°C for 3 minutes. Amplification was confirmed with gel electrophoresis using a 1% agarose gel. DNA sequencing was performed at the ISU DNA Facility with an automated DNA sequencer (ABI 3130XL; Applied Biosystems Instrument, Carlsbad, CA, USA). The results blasted with NCBI GenBank database for species assignment.

### Whole-genome sequencing

Two separate genomic DNA libraries were prepared according to the requirements of the Illumina and Oxford Nanopore systems. A combination of long-read Nanopore MinION and short-read Illumina MiSeq platforms was used to generate the complete genome sequence of the *P. hiranonis BAI-17* strain. For Illumina sequencing, the extracted genomic DNA was fragmented by sonication using a Covaris M220 sonicator (Covaris, Woburn, MA, USA). The trimmed DNA fragment sequences were then used to prepare a shotgun paired-end library with an average insert size of 350 bp using a TruSeq DNA Sample Prep kit (Illumina, San Diego, CA, USA). The library was sequenced on an Illumina MiSeq platform (Illumina, San Diego, CA, USA) using the 150-bp paired-end sequencing mode. Base calling of the two paired FASTQ files were done with the Illumina raw sequence read data. The quality of the raw sequence reads was assessed using FastQC v0.11.8, and Trimmomatic v0.39 was used to trim sequencing adapter reads with a quality score <20 at 3’ and 5’ ends.^69^ After trimming the adaptors and filtering low-quality reads, the clean sequence data were used for further bioinformatics analyses. For Nanopore sequencing, a MinION sequencing library was prepared using the Nanopore Ligation Sequencing Kit (SQK-LSK109; Oxford Nanopore, Oxford, UK). The library was sequenced with an R9.4.1 MinION flow cell (FlO-MIN106) for a 24 h run using MinKNOW v2.0 with the default settings. FAST5 files containing raw Nanopore signal data were base-called and converted to FASTQ format in real-time using Guppy v3.3.0, after which Porechop v0.2.4 was used to trim barcode and adapter sequences. Filtlong v0.2.0, a quality filtering tool for Nanopore reads, was used to remove sequences shorter than 3000 bases and with mean quality scores of less than 12 to facilitate assembly.

### De novo genome assembly and comparative phylogenomic analysis

Hybrid de novo genome assembly was performed using Unicycler v0.4.8, which allows for both Illumina reads (short, accurate) and Nanopore reads (long, less accurate) to be used in the conservative mode.^70^ The highly accurate Illumina reads were aligned against the Nanopore reads as a reference to correct random sequencing errors and finally create a genome assembly of high quality. Ultimately, the assembled sequence was polished by aligning the Illumina paired-end reads with the BWA-MEM algorithm using Pilon v1.2.3 for several rounds to improve genome assembly quality.^71^ Multiple rounds of error correction were performed until no more errors could be fixed. The quality of the assembled genome was assessed using Quast.^72^ Finally, the complete genome sequence of *P. hiranonis BAI-17* strain was annotated using Prokka 1.14.6.^73^

The *P. hiranonis BAI-17* strain was phylogenomically compared with other known primary bile acid converting species. To this end, seven genomes of *Clostridium scindens* (*Clostridium scindens* BL389WT3D, *Clostridium scindens* VE202-05, *Clostridium scindens* MSK.1.26, *Clostridium scindens* BL-398-WT-3D, *Clostridium scindens* MGYG-HGUT-01303, *Clostridium scindens* MSK.1.16, *Clostridium scindens* ATCC35704), four genomes of *Clostridium hylemonae* (*Clostridium hylemonae* MGYG-HGUT-01710, *Clostridium hylemonae* DSM15053, *Clostridium hylemonae* DSM-15053, *Clostridium hylemonae* BSD2780061688st1_A6, *Clostridium hylemonae* BSD2780061688b-171218_A6) and one human origin *P. hiranonis* genome (*P. hiranonis* DSM 13275) were downloaded from the National Center for Biotechnology Information (NCBI) and compared against the genome of *P. hiranonis BAI-17* strain using Anvio version 6.2.^74^ Briefly, first, an ’Anvi’o genomes storage database’ was generated from the FASTA files of 14 isolate genomes from *P. hiranonis BAI-17*, *C. scindens*, and *C. hylemonae* using the ’--external-genomes’ flag and Anvi-pan-genome’ with the genomes storage database, the flag ’--use-NCBI-blast,’ parameters ’--minbit 0.5′, and ’--mcl-inflation 10′ were then employed to calculate similarities of each amino acid within each genome, and remove weak, originally described in ITEP, to filters out weak hits based on the aligned fraction between two reads.^75^^76^ The MCL algorithm identified gene clusters in the remaining Blastp search results. Hence, it computed the occurrence of gene clusters across genomes and the total number of genes they contain. This generated a hierarchical clustering analysis for gene clusters (based on their distribution across genomes) and genomes (based on gene clusters they share) using a default Euclidean distance and Ward clustering. The final product was a generated anvi’o pan database that stored all results for downstream analyses to be visualized by the ’anvi-display-pan program.^77^ ’ Anvi’o also contains anvi-compute-genome-similarity, a program that uses various similarity metrics, such as PyANI, to compute average nucleotide identity (ANI) across genomes, and Sourmash to compute mash distance across genomes. The GFF3 file containing both sequences and annotations was used to compare and identify candidate genes based on Venn diagrams.^78^

### Phenotype microarrays

The *P. hiranonis BAI-17* strain was grown overnight in a pre-reduced BHI broth medium supplemented with 0.3% sodium taurocholate inside the anaerobic chamber (Coy Laboratories, Grass Lake, MI, USA). Approximately 200 µl of the overnight culture was anaerobically plated on prereduced BHI agar supplemented with 0.3% sodium taurocholate. The agar plates were anaerobically incubated at 37°C for 72 h (to ensure a thick layer of growth of bacteria). Then, sterile cotton swabs were used to pick the bacterial culture from the plate without reaching the agar plate surface and mixed with the anaerobic BIOLOG buffer. A baseline OD630 value of ∼0.02 for the buffer was ensured before substrate OD measurement. One hundred microliters of the OD630 ∼0.02 buffer was pipetted into each well of the BIOLOG PM1-PM8 plates in quadruplets. The OD630 for the bacterial and buffer mixture was measured using a flat bottom untreated 96 well plate with the Bio-tek Microplate reader (Elx808). OD630 readings were taken at 0 h and 24 h. Final OD values were divided by initial OD values to obtain fold changes in growth. The mean of the four replicates was calculated and normalized by mean fold change in the negative control to calculate % bacterial growth in each substrate. Twenty percent or greater growth in the substrate, when compared to control, was considered as significant growth.

### Bile acid transformation assays

All bile acid-containing media and controls (without bile acid) were inoculated with 200 μL of 18 h old *P. hiranonis BAI-17* strain culture. Freshly prepared stock solutions of all bile acids (CA, CDCA and TCA) were prepared using solvents (ethanol for CA and CDCA and water for TCA) and subsequently added to 20 mL BHI broth to reach a final concentration of 50 µM for the respective bile acid. The tubes were incubated in the anaerobic workstation (AS-580, Anaerobe Systems, Morgan Hill, CA, USA) at 37°C for 72 h. A sterile control and an additional control without bile acid (but with the ethanol vehicle) were included in triplicates for this experiment. Culture samples were drawn at 0h, 12h, 24h, 48h, and 72h, and the OD600 was measured soon afterward. Each sample (1 ml) was centrifuged at 5,000 x g at 4°C for 15 minutes. The supernatant was collected and aliquoted to multiple microcentrifuge tubes. All samples were stored at

−20°C until bile acid analysis (described later in this section).

### Coculture experiments, bacterial growth and toxin quantitation

Mono and Co-culture of *C. difficile* UK1 strain and *P. hiranonis BAI-17* strain were done in BHIS medium (Brain Heart Infusion supplemented with 5 g/L yeast extract) with three concentrations of CA (0, 50, and 150 μM).^79, 80^ Cocultures, and the corresponding monocultures, were incubated in triplicate under anaerobic conditions at 37°C for 48 h. Samples were drawn at 0 h, 24 h, and 48 h for the bacterial growth assessment by dilution and plating on pre-reduced blood agar and total toxin quantitation by ELISA. Each sample (1 mL) of each monoculture and coculture was centrifuged at 5,000 × g for 15 min, and pellets and supernatants were collected and stored at -20°C.

The supernatant was diluted and was assayed for *C. difficile* toxin A/B ELISA using Techlab Tox A/B kit (Techlab, Blacksburg, VA, USA) per the manufacturer’s instructions. Quantification of the *C. difficile* total toxins A and B was based on the standard curve of purified toxins generated previously.^81^ The optical density was measured at 450 nm and compared to the standard curve’s linear range to determine total toxin concentration.

### Animals, inoculum preparation, and ASF+ mouse test

All animal experiments were performed in accordance with the protocols approved by ISU Institutional Animal Care and Use Committee (IACUC-19-166, IACUC-20-091). ASF mice were obtained from the in-house breeding colonies maintained at the Vaccine Research Institute at ISU by Dr. Michael Wannemuehler. The treatment groups were housed separately and provided with irradiated feed and autoclaved water *ad libitum*. The integrity and composition of the ASF flora in the gut of the mice were confirmed by routine fecal sampling, multiplex PCR for constituent ASF flora, and 16S rRNA gene sequencing.^46^ Nine, 6-week-old (young adult) ASF mice were administered a single oral gavage dose of 100 µl of 1x10⁸ CFU/mL *P. hiranonis BAI-17* grown anaerobically in BHI medium for 24 h. Six ASF mice were kept as a control for the study. For detection of *P. hiranonis BAI-17* colonization, fresh fecal pellets were collected from individual mice at 3 and 6-weeks post-inoculation. The animals were euthanized at 3- and 6-weeks post-inoculation, and the cecal contents and intestinal tissue samples were collected for further processing.

### Real-time PCR

The fecal pellets from ASF mice were thawed, and the DNA was extracted using the QIAamp PowerFecal proDNA kit (Qiagen, Germantown, MD, USA). DNA from *C. butyricum,* a negative control, and *P. hiranonis BAI-17* positive control, were also extracted from pure cultures using the QIAamp BiOstic Bacteremia DNA kit (Qiagen, USA) according to the manufacturer’s instructions. Custom canine *P. hiranonis* specific primers and probes were designed based on the whole genome sequence information of the *P. hiranonis BAI-17*, which consists of Forward: 5’AAATAGGTGCTCAGAACATGCA3’, Reverse: 5’TCATCAGTTTCGTTGAAGTACTGT3’, and Probe: FAM-5’TGGAGCTTTCACTGGAGAAGTTGCACCA3’-TAMRA. A TaqMan-based real-time PCR was then performed on the DNA samples using the QuantStudio 3 Real-Time PCR Systems (ThermoFisher Scientific, Waltham, MA, USA), using optical grade 96-well plates. Twenty microliter reactions were made using TaqPath™ 1-Step RT-qPCR Master Mix (ThermoFisher Scientific, Waltham, MA, USA), with 10 µmol 1:1:1 ratio of forward/reverse primer, and probe and 20ng of DNA template for each reaction. The thermocycler conditions for real-time PCR were 95°C for 2 min, 95°C for 3 seconds for 40 cycles, and 60°C for 30 seconds. Each sample was run in triplicate, and the mean values were calculated. In addition, an amplification plot was generated with the threshold cycle (CT) values, and automatic analysis settings determined baseline settings.

### RNAscope® *in situ* hybridization and histopathology

Gut colonization of *P. hiranonis BAI-17* in ASF mice (ASF^+^)was confirmed by RNA *in situ* hybridization using RNAscope®, described previously.^82^ Briefly, custom paired B-*P. hiranonis*-16S (cat. 1057061-C1) double-Z oligonucleotide probes were designed against target RNA (16s rRNA) and probed region (ZZ probe targeting 17445 to 17482 nt), using custom software. The RNAscope® 2.5 HD Red Reagent (Advanced Cell Diagnostics, Newark, CA) was used according to the manufacturer’s instructions. Formalin-fixed paraffin-embedded (FFPE) tissue sections of 5 μm thickness were prepared according to the manufacturer’s recommendations. Each sample was quality controlled for RNA integrity with a probe specific to the housekeeping gene. To achieve interpretable results, assays using archived FFPE tissues were run parallel with positive and negative controls. Negative control background staining was evaluated using a probe specific to the bacterial *dapB* gene. Brightfield/Fluorescent images were acquired using an Olympus Bx53 microscope (Olympus Optical Company, Tokyo, Japan) under 40x objective.

The colon and cecum were sectioned at 5μm thickness and stained with hematoxylin and eosin. Images of these sections were acquired with 100× magnification using an Olympus BX53 microscope (Olympus Optical Company, Tokyo, Japan). The sections were evaluated by a board-certified pathologist.

## Targeted metabolomic analysis

The targeted metabolomics analysis of fecal bile acids was performed by Pennchop Microbiome and Metabolomics Core (Philadelphia, PA, USA) as described previously.^83^ A Waters Acquity uPLC System with a QDa single quadrupole mass detector was used to measure bile acids. Fecal samples were suspended in methanol (5 μL/mg stool), vortexed for 1 minute, then centrifuged twice for 5 minutes at 13,000 × g. On an Acquity uPLC with a Cortecs UPLC C-18 + 1.6 mm 2.1× 50 mm column, the supernatants were examined. The flow rate was 0.8 mL/min, and the injection volume was 4 μL, the column temperature was 30 °C, the sample temperature was 4 °C, with a run time of 4 minutes per sample. Eluent A was 0.1% formic acid in water; eluent B was 0.1% formic acid in acetonitrile; the weak needle wash was 0.1% formic acid in water; the strong needle wash was 0.1% formic acid in acetonitrile. The seal wash was 10% acetonitrile in water. The gradient was 70% eluent A for 2.5 min, 100% eluent B for 0.6 min, and then 70% eluent A for 0.9 min. The mass detection channels were: +357.35 for chenodeoxycholic acid and deoxycholic acid; +359.25 for lithocholic acid; −407.5 for cholic, alphamuricholic, betamuricholic, gamma muricholic, and omegamuricholic acids; −432.5 for glycolithocholic acid; −448.5 for glycochenodeoxycholic and glycodeoxycholic acids; −464.5 for glycocholic acid; −482.5 for taurolithocholic acid; −498.5 for taurochenodeoxycholic and taurodeoxycholic acids; and −514.4 for taurocholic acid. Samples were quantified against standard curves of at least a five points run, performed in triplicate. Standard curves were done at the beginning and end of each metabolomics run. Quality control checks (blanks and standards) were run every eight samples. Results were rejected if the standards deviated by greater than ±5%.

### Statistics

Statistical analysis was performed using GraphPad Prism 9 (GraphPad Software, San Diego, CA). The results of the animal study were expressed as means ± standard errors of the means (SEM). The differences between the experimental groups were compared using the analysis of variance (ANOVA). The differences between the two groups were analyzed using an unpaired Student’s t-test. The statistical significance level was set at p < 0.05.

### Availability of data

The genome data will be available at GenBank (https://www.ncbi.nlm.nih.gov/genbank/) following a 12-month embargo from the date of publication to allow for commercialization of research findings. The data supporting the findings of this study are available from the corresponding author SM on request after the embargo period.

## Supplementary data

**Supplementary figure 1:**
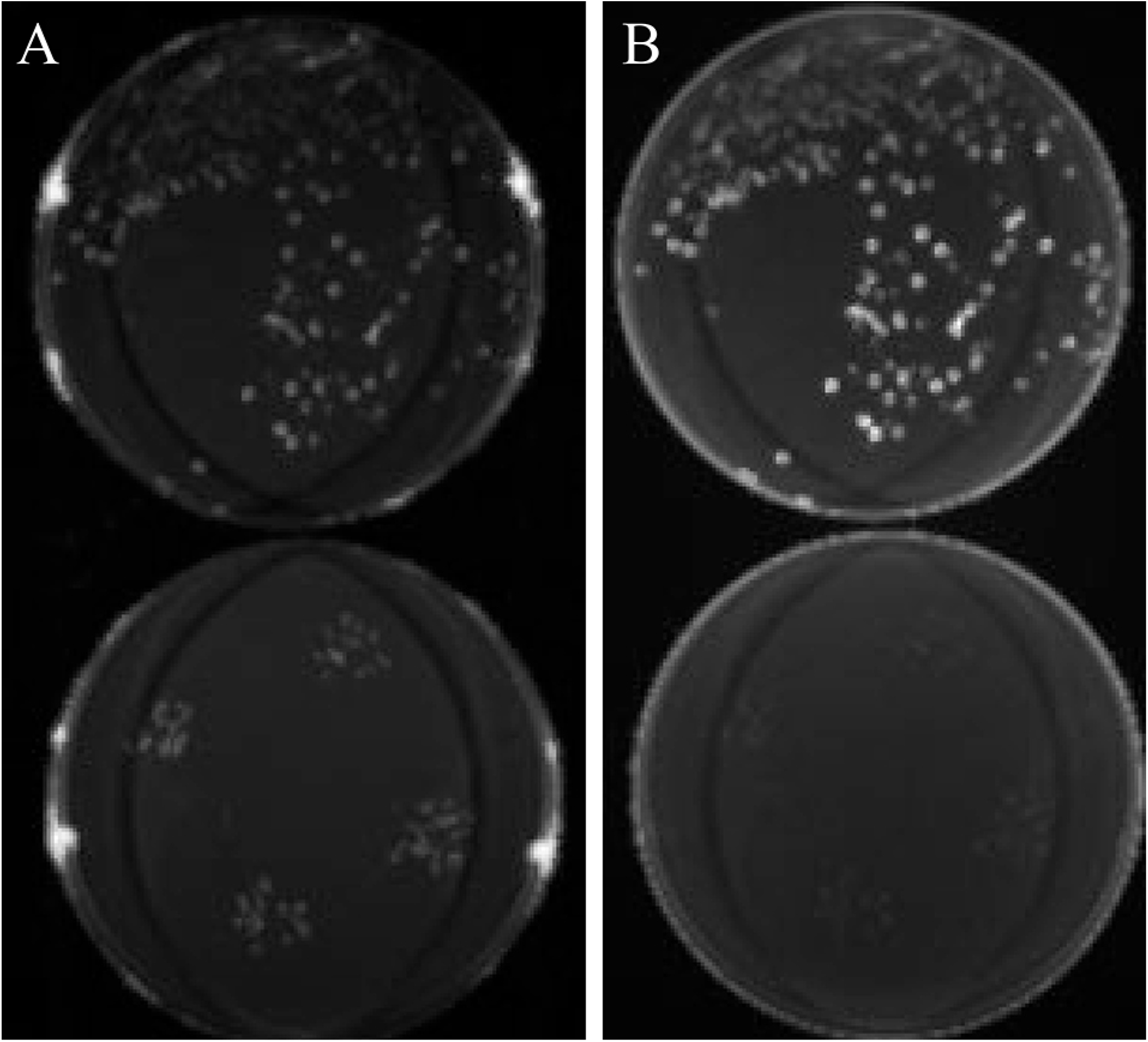
Fluorescence under ultraviolet (UV-6) discriminates *C. difficile* colonies from *P. hiranonis* colonies on blood agar. *C. difficile* UK1 (petri dish on the top in panels A and B) and *P. hiranonis* BAI-17 (petri dish on the bottom in panels A and B) were cultured on blood agar and incubated in anaerobic conditions for 48 h at 37°C. The colonies were visualized under visible light (panel A) and UV-6 (panel B) using a G BOX chemi XRQ UV transilluminator (Syngene, Frederick, MD). *C. difficile* colonies were brightly illuminated (top petri dish, panel B) under UV, while *P. hiranonis* colonies were illuminated only under visible light (bottom petri dish, panel B).

**Supplementary Table 1.** Bacterial species isolated from 270 random canine fecal samples using CCFA medium. Two seventy random canine fecal samples were enriched in BHIS medium in anaerobic conditions for 48 h at 37°C for seven days, followed by three days of oxygen shock. 1 ml of the media was centrifuged, the pellet was alcohol shocked with 100% ethanol for 20 minutes and plated on a CCFA agar plate supplemented with 0.1% taurocholate. The identity of Pro DISK positive colonies was confirmed by MALDI-TOF and 16s rRNA sequencing. * Indicates *bai*-containing bacteria.

| Bacteria | Isolation rate (%) |
| --- | --- |
| *C. baratii* | 0.37 |
| *C. boltae* | 0.37 |
| *C. butyricum* | 0.37 |
| *C. celerecrescens* | 0.74 |
| *C. cochlearium* | 2.59 |
| *C. colicanis* | 0.37 |
| *C. difficile* | 23.70 |
| *C. innocum* | 1.85 |
| *C. paraputrificum* | 2.22 |
| *C. perferingens* | 2.59 |
| *C. ramosum* | 0.37 |
| *C. sordelli* * | 0.37 |
| *C. sphenoides* | 0.74 |
| *C. sporogenes* | 5.93 |
| *C. tertium* | 4.07 |
| *P. bifermentans* * | 4.44 |
| *T. glycolicus* | 4.44 |

